# Long-lasting salt bridges provide the anchoring mechanism of oncogenic KRas-4B proteins at cell membranes

**DOI:** 10.1101/2020.08.14.250738

**Authors:** Huixia Lu, Jordi Martí

## Abstract

Ras is a family of related proteins participating in all animal cell lineages and organs. Ras proteins work as GDP-GTP binary switches and regulate cytoplasmic signalling networks that are able to control several cellular processes, playing an essential role in signal transduction pathways involved in cell growth, differentiation and survival so that overacting Ras signalling can lead to cancer. One of the hardest challenges to face is, with more than hundred different missense mutations found in cancer, the design of mutation-selective therapeutic strategies. In this work, a G12D mutated farnesylated GTP bound KRas-4B protein has been simulated at the interface of a DOPC/DOPS/cholesterol model anionic cell membrane at the all-atom level. A specific long-lasting salt bridge connection between farnesyl and the hypervariable region of the protein has been identified as the main mechanism responsible of the binding of oncogenic farnesylated KRas-4B to the cell membrane, since this particular bond is absent in both wild-type and oncogenic methylated species of KRas-4B. This finding may lead to a deeper understanding of the mechanisms of protein binding and eventual growing and spreading inside cell membranes. From free energy landscapes obtained by well-tempered metadynamics simulations, we have characterised local and global minima of KRas-4B binding to the cell membrane revealing the main pathways between anchored and released states.

## 1 Introduction

Cell membranes are active agents in the regulation of protein structure and function, including those involved in cancer diseases such as the proteins belonging to the Ras family^1^. In particular, specific mutations in the family of Ras proteins are known to be responsible of their conversion to oncogenic species^2,3^. Ras proteins family members are small guanosine-triphosphatase (GTPase) species that can be activated or de-activated by guanosine-diphosphate (GDP)-guanosine-triphosphate (GTP) switching, what means that in most cases Ras proteins oscillate between inactive GDP-bound and active GTP-bound states to regulate cell survival and proliferation^4^. As a surface protein anchored in the inner leaflet of the cell membrane (i.e. at the cytoplasm), Ras proteins are normally in the inactive state. They are activated following an incoming signal from their upstream regulators. The regulation between GDP and GTP is usually performed by loss of GAP (GTPase Activating Proteins, like neurofibromin) or by activation of GEF (Guanine Exchange Factors, like the human regulator of chromosome condensation, RCC1). Ras proteins play an essential role in signal transduction pathways which ultimately turn on genes involved in cell growth, differentiation and survival, eventually leading to cancer^5^.

Conventionally, mutant Ras is considered to be defective in GAP-mediated GTP hydrolysis, which results in an accumulation of constitutive GTP-bound RAS in cells. Missense gain-of-function mutations in all three Ras genes are found in 27% of all human cancers, with 98% of the mutations at one of three mutational hotspots: G12, G13 and Q61. Historically, the majority of biochemical and structural studies of Ras have focused on HRas^6^. However, HRas is the least frequently mutated Ras isoform in human cancers (4%), whereas KRas is the predominantly mutated isoform (85%), followed by NRas (11%). It has been observed that it was KRas G12D mutation instead of NRas G12D mutation which promoted colon cancer development in Apc-deficient mice, supporting the ability of KRas but not NRas to initiate the formation of such cancers^7^. Consequently, we have chosen the specific mutated G12D KRas-4B as our target oncogenic protein.

KRas has two splice variants, KRas-4A and KRas-4B, the latter being found at higher concentration levels^8^, in particular in lung, colorectal, and pancreatic cancer cells^9–11^. These insights suggests the importance of fully understanding the regulation of oncogenic KRas-4B activity when binding on the membrane. KRas-4B has a highly conserved N-terminal catalytic domain (CD, residues 1-166) and a flexible C-terminal 22-25-amino acid-long, hypervariable region (HVR, residues 167-185)^12^. Apart from different HVR sequences, KRas-4B is distinguished by its unique HVR post-translational modification (PTM). Usually, HVR lipidation promote Ras anchoring in the plasma membrane. Furthermore, unlike other Ras isoforms, the HVR of KRas-4B is polybasic. HVR preferentially binds the membrane in the liquid phase and spontaneously inserts its terminal farnesyl (FAR) moiety into the loosely packed phospholipid bilayers^8^.

Accumulating evidence indicates that demethylated and farnesylated KRas-4B (KRas-4B-Far, see Fig. 1 of “Supplementary Information”, SI) could play an important role in the signalling pathway that happens on the inner leaflet of the membrane bilayers. KRas-4B-Far has been reported to be able to be transferred to bind the inner plasma membrane (PM) leaflet. According to Ntai et al. 91% of the mutant KRas-4B and 51% of wild-type KRas-4B proteins in certain colorectal tumor samples have been found to be of the KRas-4B-Far isoform^13^. Nevertheless, the effects of KRas-4B-Far on downstream signalling have yet to be determined. While most efforts have been focused on characterisation of methylated KRas-4B (KRas-4B-FMe) binding to the PM, study on KRas-4B-Far is still an emerging area of research. According to Barcelo et al.^14^, phosphorylation at Ser-181 of oncogenic KRas is required for tumor growth so that in the present work we have chosen to phosphorylate at Ser-181 to model the main mutated oncogenic KRas-4B protein under study.

**Figure 1:**
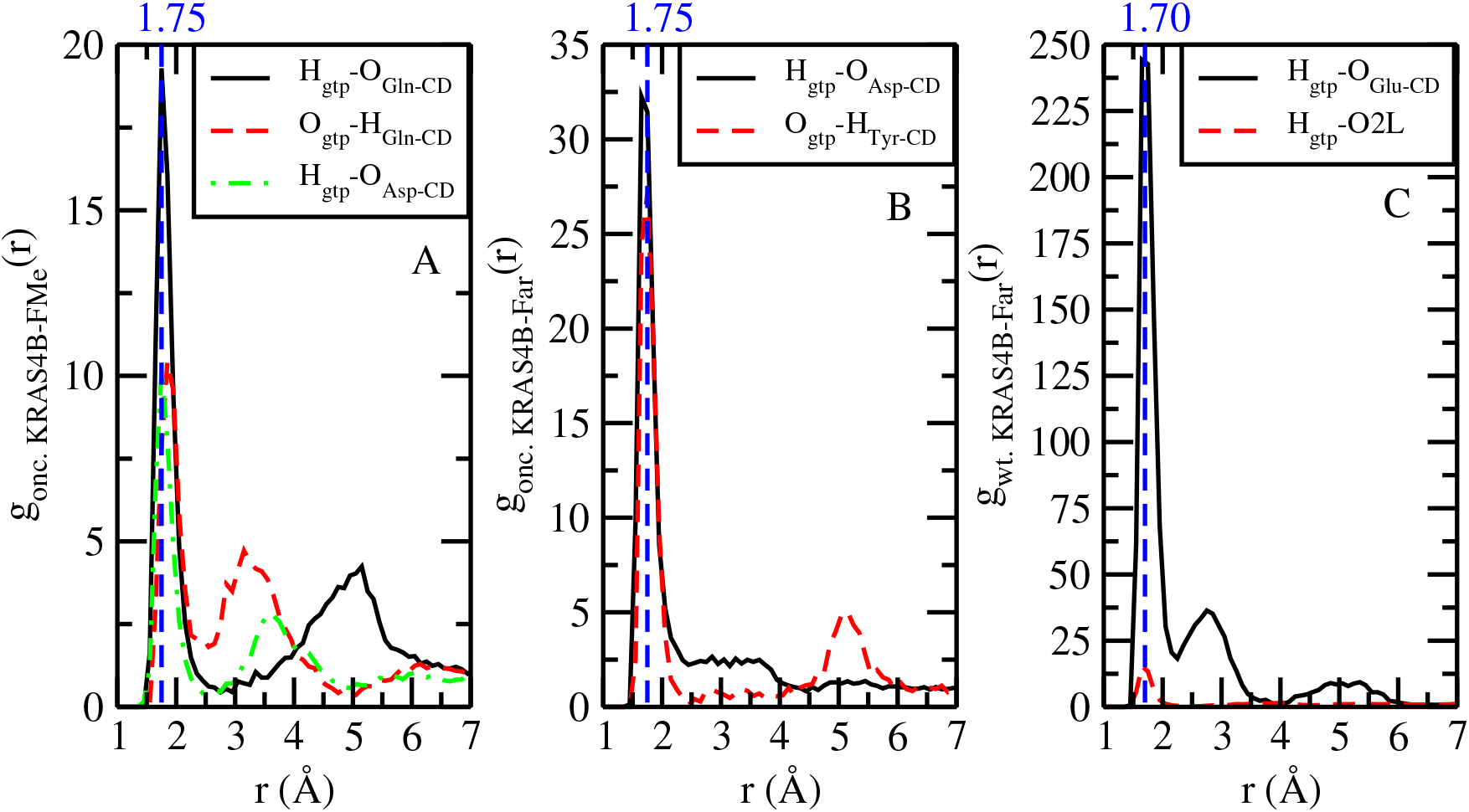
Selected RDF for active atoms of GTP with selected sites of the CD and active oxygen atoms of head groups of lipids (’O2L’). ‘H_*gtp*_’ and ‘O_*gtp*_’ represent hydrogen and oxygen atoms from phosphate group of GTP.

In particular we shall provide essential information in order to gain precise understanding of the effects of oncogenic mutations on the localisation of wild-type and oncogenic KRas-4B proteins and on their eventual mechanisms to anchor the cell. According to Nussinov et al.^15^, the two major pathways in oncogenic Ras-driven proliferation can be promoted when KRas is membrane-anchored. Furthermore, understanding the structural specifics of KRas-4B in its GTP-bound form will help to design oncogenic KRas-4B inhibitors^16^. So, methods of reducing or weakening the interactions between oncogenic KRas-4B and the membrane with a good knowledge of the structural mechanisms at the atomic level will be a promising target for anticancer drug discovery^17^.

On its own, genomics sequence data may not provide the entire information to the oncologist for the selection of targets. Further, the GTP affinity of KRas-4B is reported to be extremely high, with a dissociation constant of around 10 mol/L ^18^, yet the corresponding binding free energy hypersurface has been poorly explored. The free energy landscape (FEL) idea is compelling because it enables us to map many possible conformations which the protein could populate along the different levels of free energies so that recommendations for treatments might derive from the corresponding FEL, producing further insight into the underlying biological mechanisms and fostering molecular targeting in a significant way^19^. Ras association with membranes is not a one-way street and they undergo a cycle of delivery to the PM followed by return to endomembranes for recycling^20^. In this work we have applied well-tempered metadynamics to reveal the affinity of GTP to KRas-4B and lipids and to obtain the binding free energy barriers of the process of anchoring of FAR at membrane bilayers from a free energy perspective, especially for membranes with relatively high contents of cholesterol. For the first time, detailed free energy landscapes of FAR and GTP for different KRas-4B sequences are reported.

## 2 Results

### Structure of KRas-4B cell membranes

The main PTM transformations and the three sequences of farnesylated KRas-4B proteins considered in this work are shown in Fig. 1 of SI. There we compare the wild-type KRas-4B-Far protein with two modified species in which we applied mutation G12D at Gly-12 and phosphorylation (PHOS) at the Ser-181 site, namely species KRas-4B-Far and KRas-4B-FMe. Sketches of parts of the full systems are described in Fig. 2 of SI.

**Figure 2:**
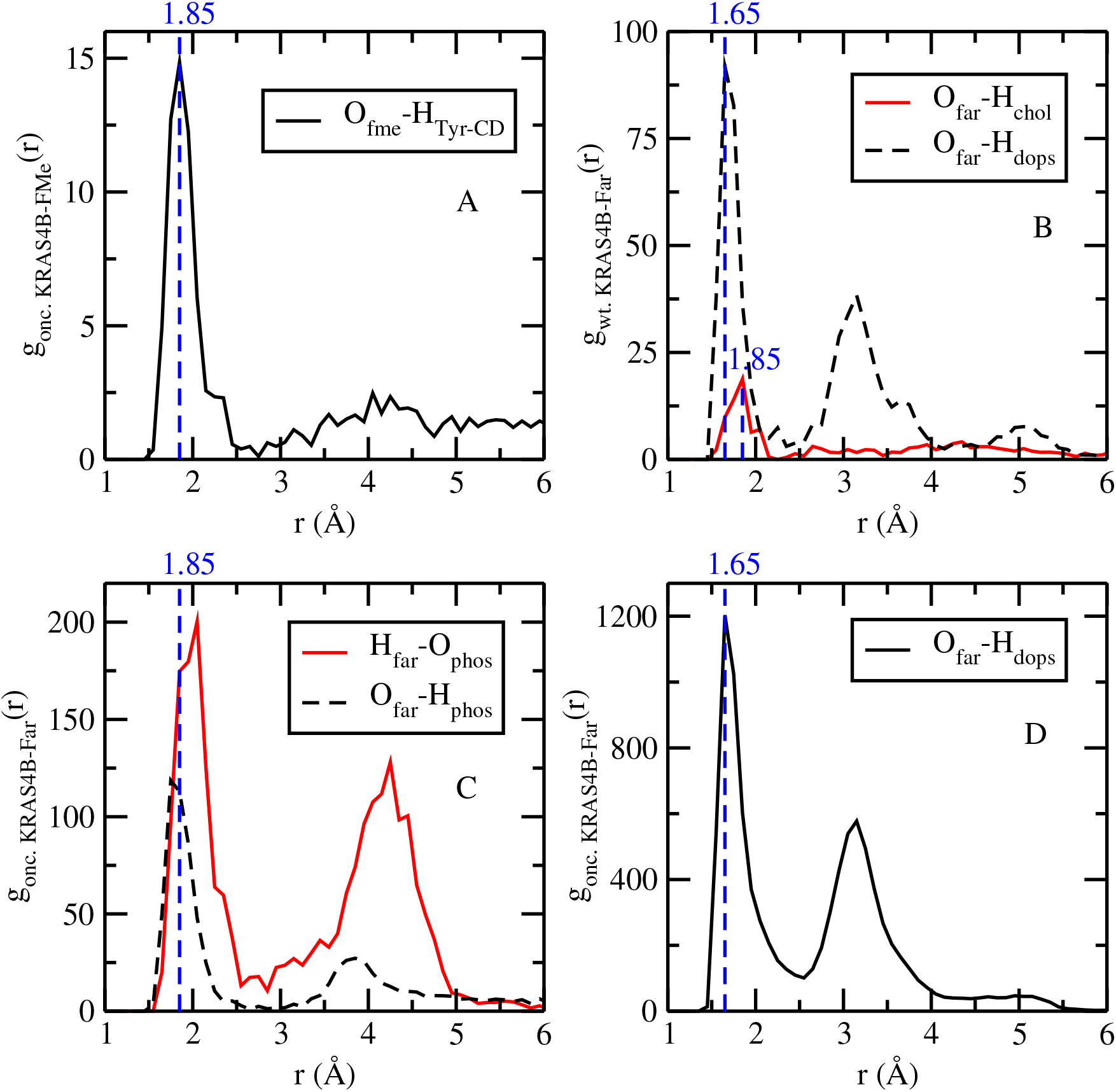
Selected RDF related to selected sites of FAR. Here O_*far*_ and H_*far*_ correspond to atomic sites belonging to farnesylated KRas-4B, whereas O_*fme*_ stands for oxygen atoms of methylated KRas-4B.

In order to explore structural characteristics of the anchoring of KRas-4B proteins at anionic membranes, several physical properties of the membrane have been calculated. A deuterium order parameter *S_CD_* is usually defined for each CH_2_ and CH groups of the DOPC lipid tails^21,22^ and averaged results are shown in Fig. 3 of SI for both tail chains of all DOPC lipids in all KRas-4B systems studied in this work. Order parameters of the first half of the chain of DOPC (carbon units 2-10) are directly related to the area per lipid, whereas the length of the whole chain is directly related to the thickness of the membrane. By comparing the present *S_CD_* with results of pure DOPC bilayers, we find a good agreement with results provided in Refs.^23,24^ in all cases.

**Figure 3:**
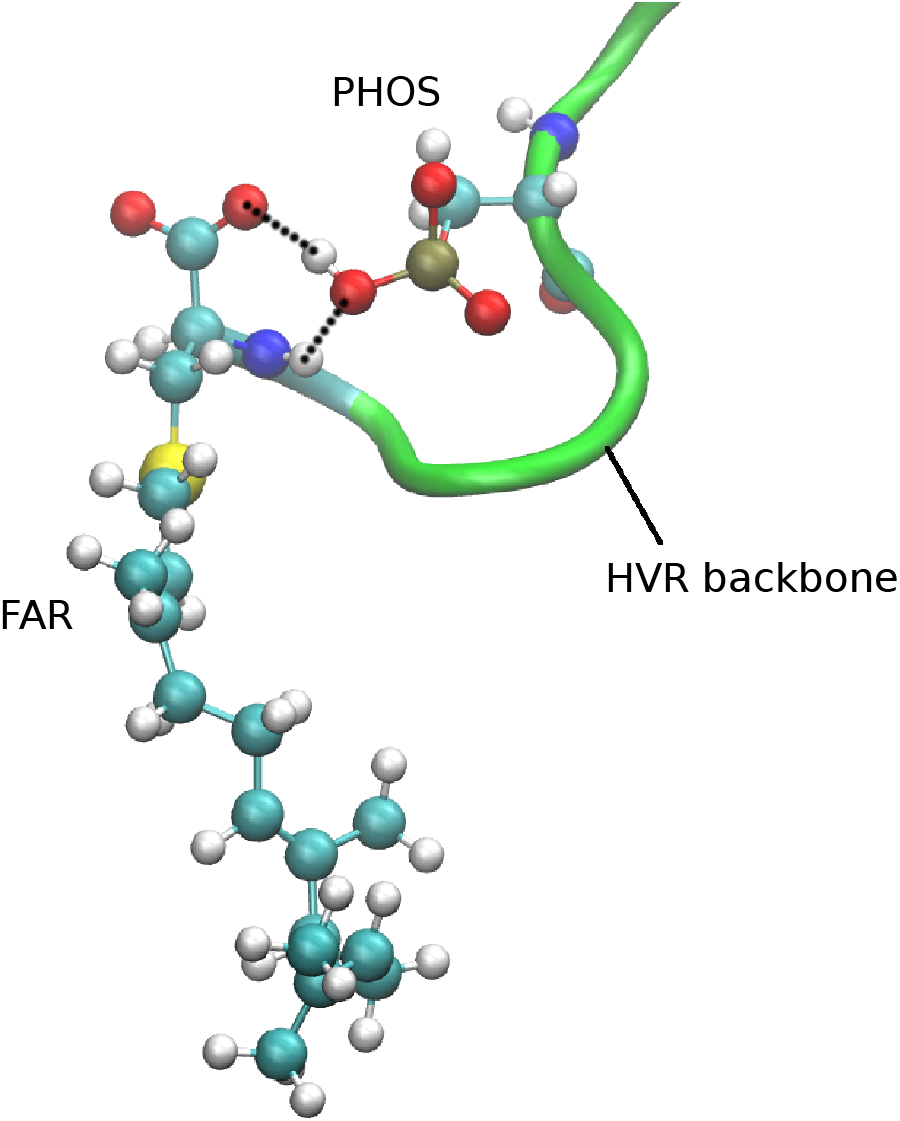
Snapshot of long-lasting salt bridges indicated as black lines of dashes between FAR and PHOS of the oncogenic KRas-4B-Far. The atoms represented here are: carbon (cyan), oxygen (red), hydrogen (white), nitrogen (blue) and phosphorus (brown). Hydrogen bonds are represented in black dashed lines.

Given the addition of 30% cholesterol, we observe that at 1 atm and 310.15 K the liquid phase was safely reached in all cases since, given the transition temperatures of DOPC (*T_m_* = 253.15 K) and DOPS (*T_m_* = 262.15 K)^25^, the extent of ordering reported by *S_CD_* is very low (between 0.1 and 0.2). Area per lipid is often used as the key parameter when assessing the validity of MD simulations of cell membranes. It has been proposed that a good test for such validation is the comparison of the area per lipid and thickness of the membrane with experimental data obtained from scattering density profiles^26^. For continuous MD production runs the area per lipid as a function of simulation time has been computed and the averaged values are reported in Table 2 of SI. Obviously different sequences of KRas-4B do not have much influence on the area per lipid of the membrane. As obtained from analysis of X-ray scattering in the low angle and wide angle regions at 303 K, area per lipid of the DOPC:cholesterol (30%) bilayer was reported by Nagle et al.^27^ with a value of 0.544 nm^2^ which is close to the value we are reporting here for all systems. DOPS bilayers were found to be thicker than DOPC bilayers with a correspondingly smaller cross-sectional area which gives smaller value of area per lipid, suggesting that DOPS has a condensing effect on DOPC bilayers^28^. We obtained a value of 0.52 nm^2^ which is 26.7% less than the experimental value of 0.71 nm^2^ for area per lipid of pure mixed DOPC/DOPS (4:1) bilayers at 297 K^29^ but that is very consistent with the experimental analysis of Nagle et al. reported above^27^.

The thickness (Δ*z*) of the membrane may provide additional clues about its mechanical properties, such as its rigidity or its capability of anchoring the FAR moiety. Δ*z* has been obtained by computing the mean distance between phosphorus atoms of the DOPC head groups from both leaflets. Results are reported in Table 2 of SI. There we can observe that Δ*z* of the oncogenic KRas-4B-Far system is slightly shorter than in the case of wild-type KRas-4B-Far and of the mutant KRas-4B-FMe, by around 2.5%. This could be attributed to stronger interactions between mutated KRas-4B-Far and lipid groups and to the deep insertion of FAR in the membrane, which significantly affect the shape of the membrane at the interface, as we will show below. The experimental thickness of pure DOPC/DOPS (4:1) bilayers at 303 K bilayers were reported by Novakova et al.^30^, with values of about Δ*z* = 3.94 nm, which are smaller than those reported here for the protein-bilayer systems, essentially due to the high contents of cholesterol considered in the present work.

### Radial distribution functions

We have considered a series of radial distribution functions (RDF) between selected atomic sites (as described in Figs. 2, 4 and 5 of SI). Being the most relevant, RDF of GTP and FAR are shown in Figs. 1 and 2, respectively. Others RDF are reported in SI (related to CD, Fig. 6 and to HVR, Fig. 7).

**Figure 4:**
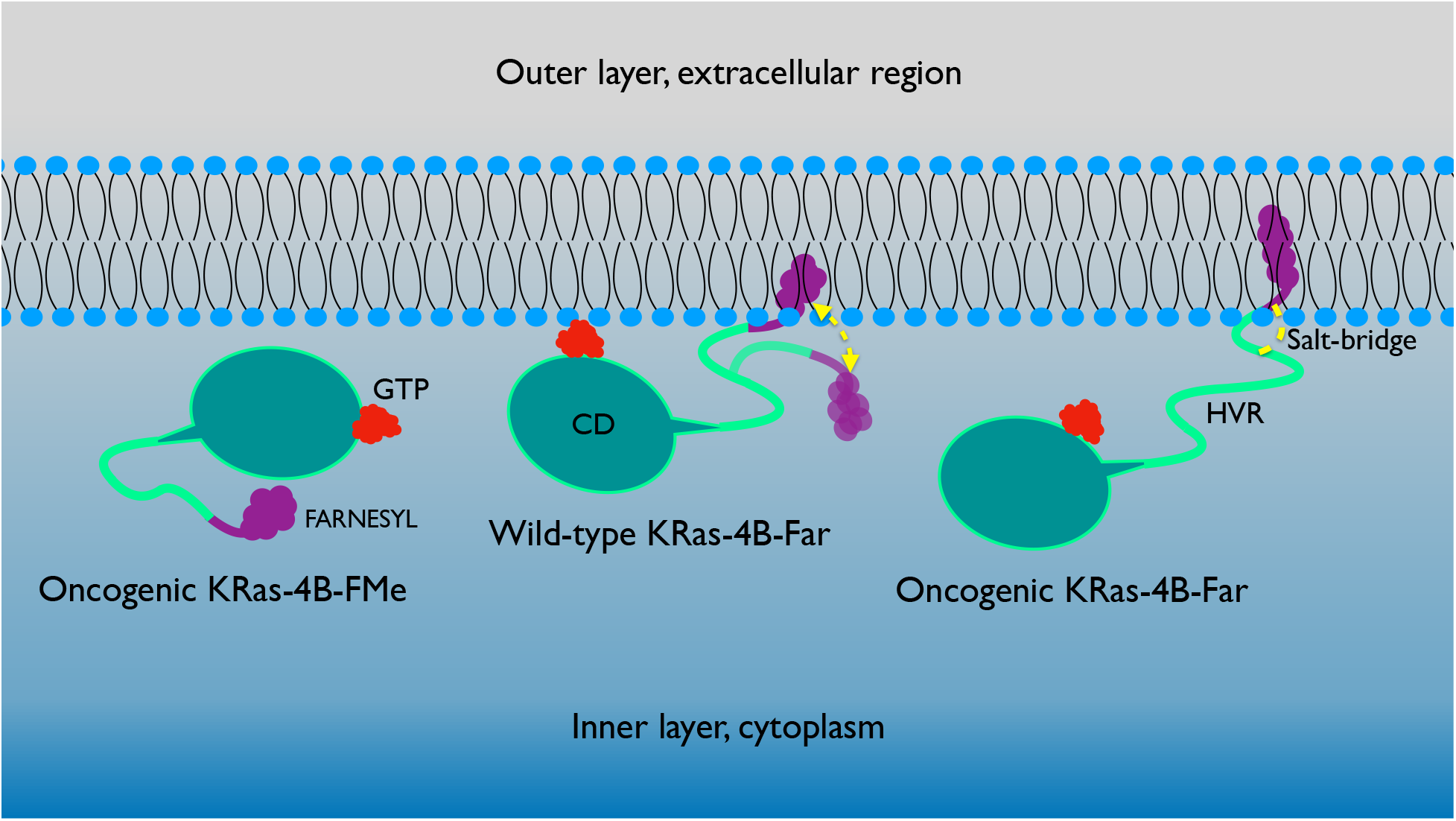
Sketch of the most stable configurations that KRas-4B-Far can adopt when attached in a cell membrane, for the wild-type and oncogenic species. Whereas wild-type can indistinctly adopt two preferential configurations (corresponding to free energy minima, as explained below) oncogenic species have only a single preferential state due to a unique free energy minimum.

**Figure 5:**
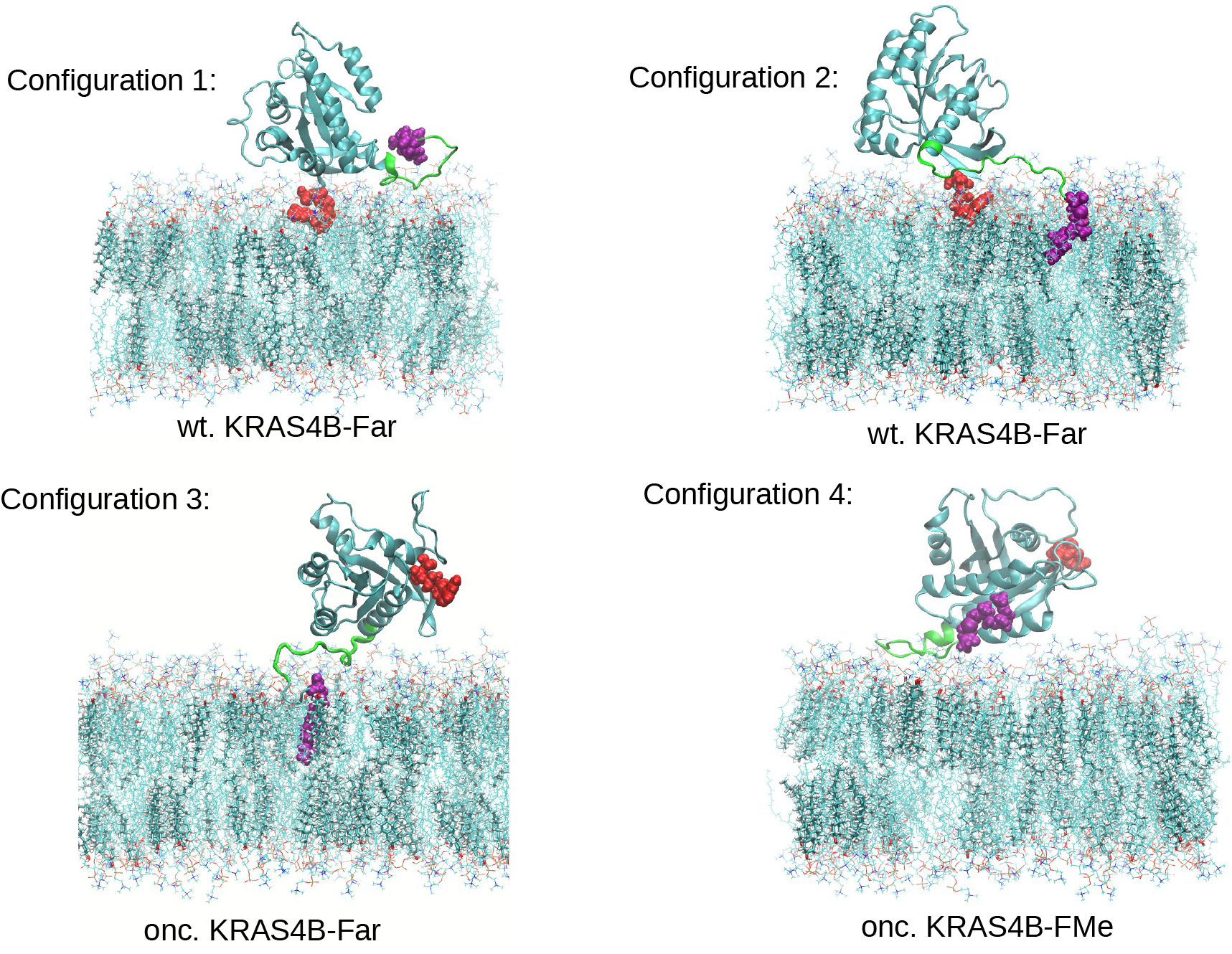
Four preferential configurations of the three KRas-4B-membrane systems obtained from MD simulations at the all-atom level.

Firstly, hydrogen-bonds (HB) between GTP and active sites of specific aminoacids of the CD have been observed. A clear first coordination shell located around 1.75 Å shall be essentially attributed to HB between selected atom sites. The typical signature of oxygen-hydrogen HB in lipid cell membranes is a maximum located at 1.8 Å^31^. For oncogenic KRas-4B systems, GTP tends to bind the CD of the protein through HB, whereas no HB between GTP and lipids have been observed. From the results reported here, we can observe that GTP prefers binding to the CD of oncogenic KRas-4B proteins regardless of carboxymethylation at site Cys-185 before anchoring to the membrane leaflet. In particular, for wild-type KRas-4B-Far, HB between GTP and the CD are much stronger (~ 15-fold) than HB between GTP and lipids. However, in the wild-type system GTP is mostly located at the interface of the membrane, close to the CD of KRas-4B. Especially strong interactions observed between GTP and CD of KRas-4B-FMe indicate less efficient nucleotide exchange for this mutant.

To further analyse the effect of the side chain of CD, calculations were performed revealing that CD is able to associate head groups of lipids, which proves that CD of KRas-4B plays a role in binding to bilayers in all cases. In the corresponding RDF (Fig. 6 of SI), when the two selected oxygen and hydrogen sites are ionised the location of the first shell is usually at distances shorter than 1.8 Å. The contribution of the first shell is from a so-called *salt bridge*^32,33^, which coupling was reported to be of general importance to the stability and function of proteins. A salt bridge has two components: a hydrogen bond and an electrostatic ionic interaction. Very common in proteins are those between the anionic carboxylate (RCOO^−^) of aspartic or glutamic acid and the cationic ammonium 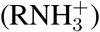 from lysine or the guanidinium 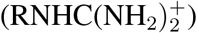 of arginine. Of all non-covalent interactions, salt bridges are among the strongest^34^. Here we report that the first shell of RDF due to salt bridges between *O_CD_* and *H_dops_* is located at 1.65 A for the three systems (the two oncogenic and the wild-type species of the protein). Nevertheless, it has been shown that HB between the cationic ammonium 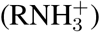 from lysine of CD of KRas-4B-FMe and anionic oxygen atoms from DOPS is much stronger than for demethylated KRas-4B isoforms.

**Figure 6:**
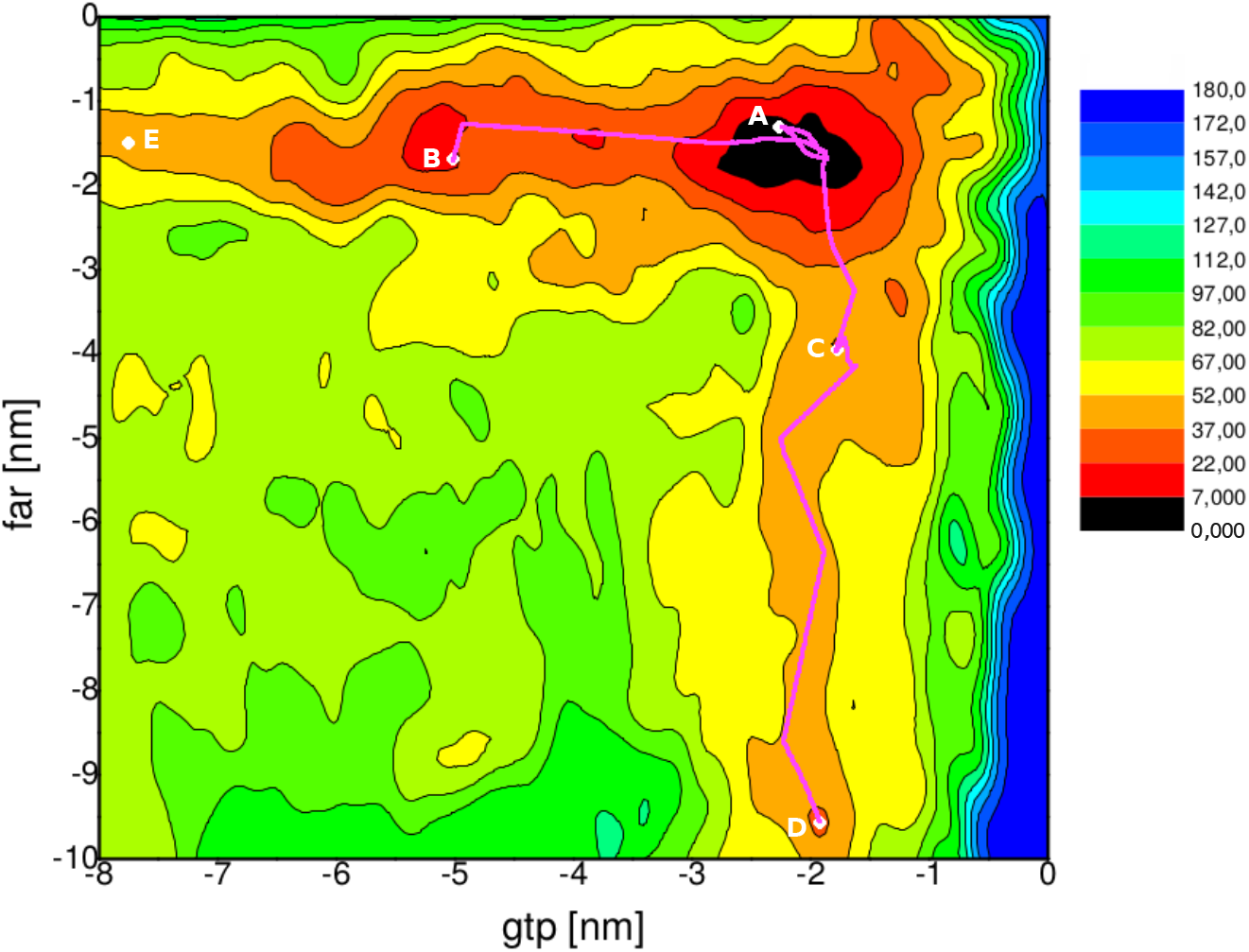
2D free energy landscapes F(gtp, far) (kJ/mol) for the oncogenic KRas-4B-Far system. Several stable and meta-stable configurations (A, B, C, D, E) are indicated. All MFEP between two selected basins are depicted in purple and the relevant coordinates have been reported in Table 4 of SI.

Consecutively, RDF related to HVR are displayed in Fig. 7 of SI. We have found that HVR is able to form HB and salt bridges with CD, membrane lipids and PHOS. Carboxylmethylation and mutations show their influence on the interactions of HVR with DOPS and PHOS (panels D and F). Whereas the first shell is located at 1.65 Å, the salt bridge between O_*HVR*_ and H_*dops*_ is much weaker for oncogenic KRas-4B-FMe, when comparing with the demethylated case. After phosphorylation (as shown in Fig. 4 of SI), oncogenic KRas-4B proteins (by means of O_*phos*_) are able to form stable salt bridges with hydrogen atoms of cationic ammonium from the HVR (H_*HVR*_). However, for wild-type KRas-4B-Far only weak HB exit between H_*HVR*_ and some oxygen atom from the side chain of site Ser-181. As it was pointed out in Refs. ^35,36^ our results suggest that CD and HVR have a significant role in binding to the PM, although engaging to the membrane in different ways.

**Figure 7:**
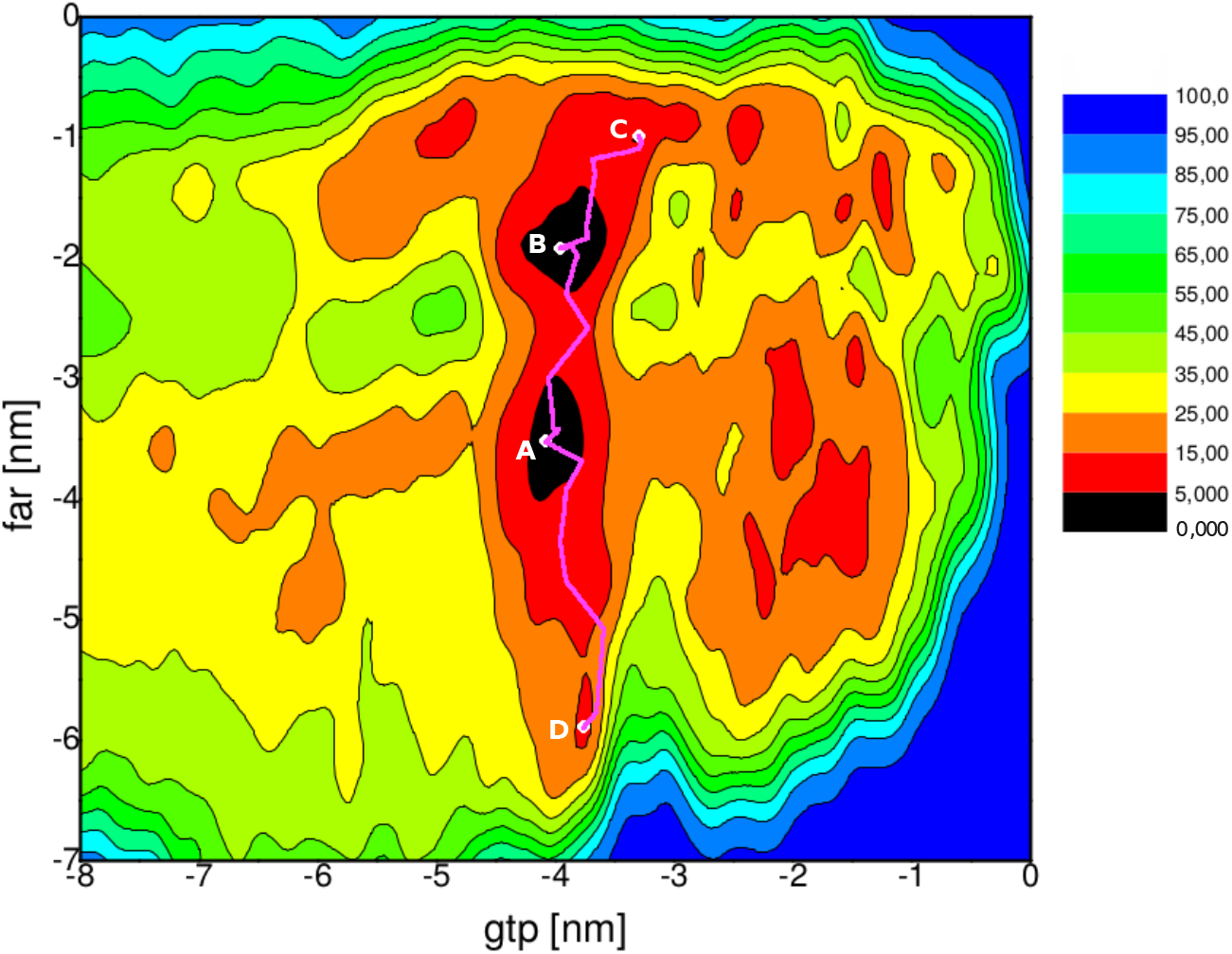
2D free energy landscapes F(gtp, far) (kJ/mol) of wild-type KRas-4B system. Stable and meta-stable configurations (A, B, C, D) are indicated. MFEP between different two basins are depicted in purple and the relevant coordinates have been reported in Table 4 of SI.

We have investigated meaningful RDF of active sites of FAR with PHOS, CD, and lipids. From Fig. 2 we obtained (panel D) that in the oncogenic KRas-4B-Far system FAR forms strong and long-lasting salt bridges with DOPS, through negatively charged *O_far_* interacting strongly with the positive charged *H_dops_* of DOPS. In the case of the oncogenic KRas-4B-FMe, HB are found (panel A) between the carbonyl oxygen of the ester group of Cys-181 (*O_fme_*), which is weakly basic, and the positive hydrogen of the hydroxyl group of Tyr from CD (H_*Tyr–CD*_). FAR can also interact with cholesterol (panel B) through typical HB and with DOPS through strong salt bridge for wild-type KRas-4B-Far, which guarantees that the GTP-bound wild-type KRas-4B-Far could anchor into anionic membrane bilayers.

However, the most remarkable fact is seen in panel C. Only in the oncogenic KRas-4B-Far long-lasting salt bridges have been located between oxygen and hydrogen atoms from FAR and PHOS, indicating that anchoring of FAR in the oncogenic KRas-4B-Far possesses a large stability provided by the permanent binding of the FAR-PHOS pair. Consequently, the anchoring mechanism described here is playing a key role in order to reach the stability required for oncogenic KRas proteins to be able to become functional, remaining attached to the inner surface of the plasma membrane through its farnesylated and poylcationic C-terminus^37^.

In the snapshot reported in Fig. 3, we described the long-lasting and strong salt bridges existing between active sites of FAR and PHOS, only for the oncogenic KRas-4B-Far protein. O1P_*phos*_ and O2P_*phos*_ could shift to form salt bridge with H_*far*_, but only one of them could interact with H_*far*_ hauling the typical HB distance, with one of them favored by H_*far*_ because of the conformational restrictions in the structure even though both of them share one negative charge. After analysing production trajectories carefully, we have found that O_*far*_, H_*phos*_, H_*far*_, O1P_*phos*_/O2P_*phos*_, the nitrogen, and two carbons of FAR form a very stable 7-membered ring endowing the oncogenic KRas-4B-Far to get such a specific structure in its tail able to allow the anchoring to PM or reacting with other proteins *in vivo*. Finally, in order to monitor the interactions between FAR and PHOS for the oncogenic KRas-4B-Far system, we displayed the time evolution of selected atom-atom distances in Fig. 8 of SI regarding the strong salt bridge observed in Fig. 2, panel C, where we see that the anchoring mechanism described here is fully stable throughout a simulation span of 500 ns. In summary, the finding or design of drugs able to break the long-lasting bonds between FAR and PHOS could be a key factor for the release of oncogenic proteins such as KRas-4B from its anchoring to the membrane towards the internal regions of the cell, eventually becoming harmless.

**Figure 8:**
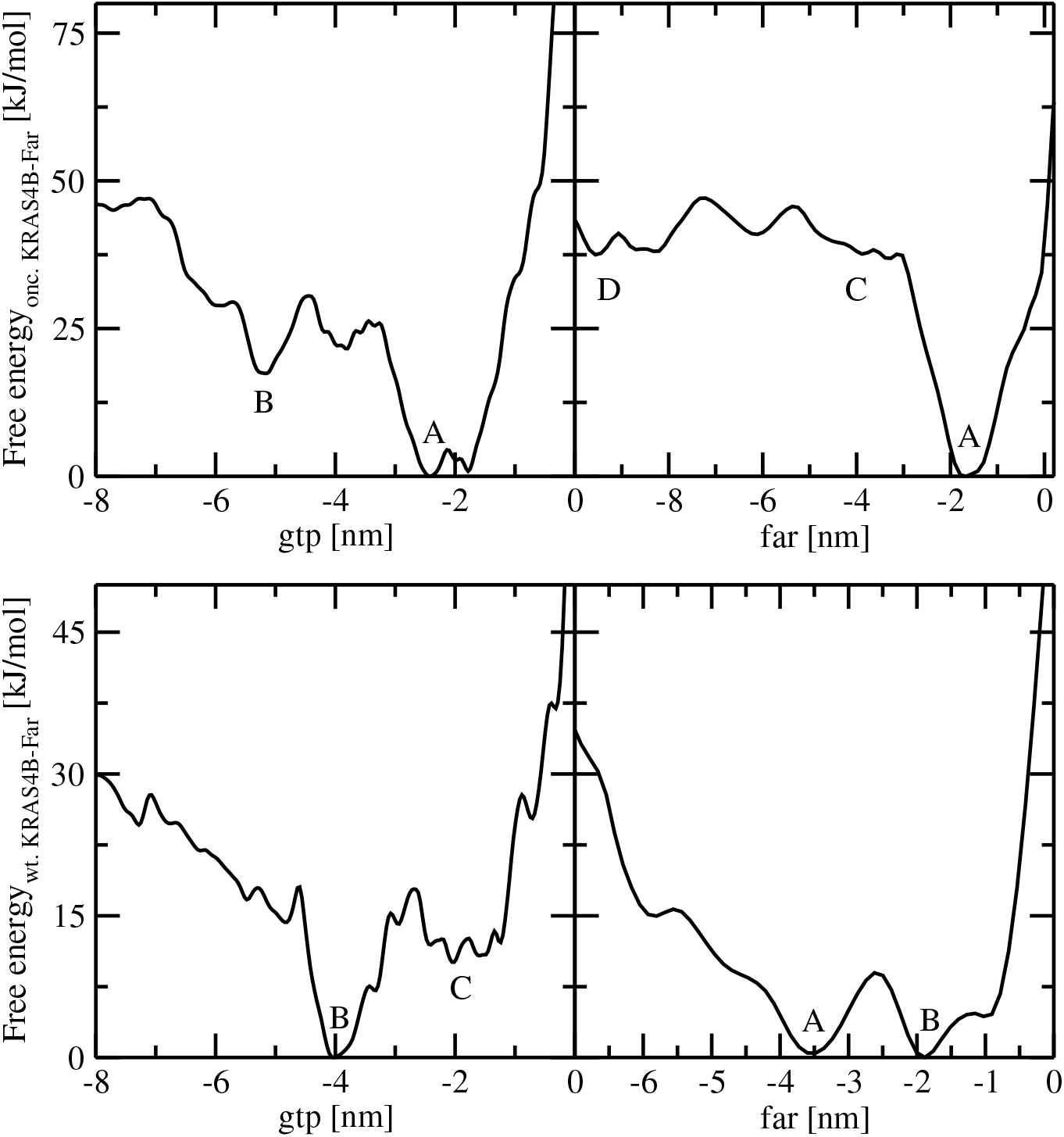
1D integrated free energy profiles of GTP and FAR along the membrane normal direction. Basins marked with same labels as in Figs. 6 and 7. Minima in each of the four panels are set equal to zero. Free energy profiles generated by the PLUMED2 tool.

### Preferential localisation of KRas-4B at cell membranes

In a real cell, where KRas is located in the inner layer of the membrane, the characteristic configurations of wild-type and oncogenic KRas-4B-Far could be sketched as in Fig. 4, according to our results. Nevertheless, at the atomic level, the four main configurations corresponding to two stable states of the wild-type KRas-4B-Far and to the main stable states of oncogenic KRas-4B-Far and KRas-4B-FMe obtained from MD are illustrated in Fig. 5 with full details. During the simulation time, tracking the movement of different domains of KRas-4B and GTP along the membrane normal gives us direct information of how the KRas-4B proteins and the GTP molecule regulate their localisation. We report in Fig. 9 of SI the Z-axis (normal to the XY instantaneous plane of the membrane) positions of center of FAR and GTP from the center of lipids (z = 0) using the second half of the 1000 ns simulations for all cases.

After thorough inspection of Fig. 9 of SI and Fig. 5, together with the averaged values reported in Table 1, we report a list of relevant findings:

1. In all cases the leaflet with KRas-4B bound to the membrane tends to be slightly thicker with lipid tails about 0.2 nm more stretched-out than in the second leaflet.
2. When wild-type KRas-4B-Far is in its configuration 1, HVR and FAR tend to wander around z = 3.90 nm in parallel along the XY plane at the interface of the membrane, whereas in configuration 2 HVR is located roughly 0.5 nm closer to the membrane center while FAR is anchored to the membrane and located around z = 1.73 nm. GTP buries itself in the interface of the membrane, with its center located around z = 2.39 nm, whereas CD is placed above the membrane XY plane with its center at position z = 4.48 nm.
3. For the oncogenic KRas-4B-Far (configuration 3), FAR is revealed to be constantly anchored to the anionic membrane. Its center is located at z = 1.37 nm with HVR located around 3.56 nm. GTP keeps bound to the CD around z = 5.16 nm, while CD is located at z = 4.7 nm.
4. In the oncogenic KRas-4B-FMe case (configuration 4), HVR interacts with lipids and the CD, with FAR flipped away from the surface of the membrane at the location of z = 3.82 nm. FAR does not insert into the membrane bilayer in any case. Instead, it is exposed and available for antibody recognition^38^. No spontaneous FAR insertion was observed during the simulations.
5. For oncogenic KRas-4B-FMe, HVR is reported to be 0.86 nm closer to the interface than the CD and it is sandwiched between the effector binding site of the CD and the membrane. Comparing to oncogenic KRas-4B-Far, the phosphate group strengthens the auto-inhibition of KRas-4B-FMe by the HVR, same as the wild-type GDP-bound KRas-4B suggested by *in vitro* and in *silico* observations^39^.
6. According to Ref.^8^, phosphorylation of Ser-181 prohibits spontaneous FAR membrane insertion for methylated KRas-4B-FMe, which is consistent with our results of oncogenic KRas-4B-FMe system. However, the prohibition effect of phosphorylation has been drastically diminished for demethylated KRas-4B-Far proteins.
7. GTP of wild-type KRas-4B system is located always at the interfacial region, while GTP is bound to the CD region of oncogenic KRas-4B proteins. From RDF reported in Fig. 1 it is revealed that in the stable state we have observed in our MD simulations, GTP serves as a direct connection between the CD and lipids through pure HB and salt bridges.

**Table 1:** Averaged values of z-locations of FAR, HVR, CD, and GTP for KRas-4B-Far system. Values of location of FAR from configurations 1 and 2 have been provided for the wild-type case. Estimated errors in parenthesis.

| Systems | FAR | GTP | HVR | CD |
| --- | --- | --- | --- | --- |
| wt. KRas-4B-Far | 3.9(0.48)/1.73(0.23) | 2.40(0.17) | 3.93(0.25) | 4.48(0.13) |
| onc. KRas-4B-Far | 1.37(0.18) | 5.16(0.60) | 3.56(0.19) | 4.70(0.28) |
| onc. KRas-4B-FMe | 3.82(0.36) | 5.07(0.94) | 3.84(0.24) | 4.70(0.41) |

In the MD simulation runs, we have observed that FAR of wild-type KRas-4B-Far is able to easily switch between configurations 1 and 2. The main reason should be attributed to the small free energy barriers between these two stable states. It is clear that for methylated KRas-4B-FMe, FAR cannot bind the membrane in the presence of PHOS, showing a blocking effect. However, since for demethylated KRas-4B-Far proteins (wild-type and mutated) PHOS favors FAR to be inserted more deeply and stably into the membrane we could have indications of the presence of a high free energy barrier avoiding the release of FAR off the membrane.

In oncogenic KRas-4B-Far system FAR is anchored deeply into the membrane while HVR is buried at the interface. HVR stays close to the interface and CD is located 0.22 nm farther to the membrane center than CD of the wild-type case. So, we have observed that CD acts much more actively, gaining more accessibility to expose its binding sites for interacting with its upstream regulators and downstream effectors. The interaction between PHOS and DOPS lipid in the oncogenic KRas-4B-Far system, such as O_*far*_ and H_*dops*_ (see Fig. 2), is formidable, almost 12-fold stronger than the same interaction from the wild-type KRas-4B-Far system, which leads FAR of oncogenic KRas-4B-Far to anchor the membrane during the whole simulation span.Our observations indicate that wild-type KRas-4B-Far can easily anchor to and be released from the membrane without crossing too high free energy barriers. However, in the oncogenic case KRas-4B-Far might face much higher free energy barriers in order to release its C-terminus tail from the anionic membranes. This eventually helped GTP-bound KRas-4B-Far always anchor to the membrane, and to remain active. In order to explore conformational states of selected reaction coordinates and understand the structural specifics of KRas-4B-Far binding to the membranes in its GTP-bound form, we have started enhanced well-tempered metadynamics simulations from the last configuration of the MD simulations and sampled the main conformational transitions of active KRas-4B along the membrane normal. The efficiency of this method has been recently proved in our lab for the case of melatonin binding to model cell membranes^40^.

### Well-tempered metadynamics simulations and conformational transitions of KRas-4B

In order to elucidate the joint effect of phosphorylation and G12D mutation on the energetic characteristics of the binding of KRas-4B in cell membranes, the FEL of wild-type and oncogenic demethylated KRas-4B-Far proteins have been computed using the well-tempered metadynamics method. Technical details are reported in Section 4 and a full study of the convergence of the simulations is reported in SI.

### Two-dimensional free energy landscapes

Free energy landscapes are dynamic and conformational states can change in response to intra- and extramolecular events^41,42^, such as the localisations of FAR and GTP. Given the specific interest of the oncogenic and wild-type KRas-4B-Far proteins in the anchoring of Ras proteins in cell membranes, we have computed two-dimensional (2D) FEL for the corresponding two sets of proteins bound to the anionic membrane composed of DOPC:DOPS:cholesterol (28:7:15) considered above. The hypersurfaces are reported in Figs. 6 and 7 and each (meta-)stable state can be indexed by a pair of CV. Several regions with clear minima are present in the two FEL. Since the range of CV space represented in Figs. 6 and 7 is rather wide, we will focus especially in the characterisation of free energy barriers between the particular states of localisations of FAR and GTP in the FEL in the region close to one leaflet of the membrane.

Since the present 2D FEL include a wide variety of (meta-)stable states, methods able to trace the minimum free energy path (MFEP) between stable states with high accuracy are highly needed. MFEP can be determined by iteratively refine a pathway connecting stable states that converges to the minimum free energy trajectories between them. MFEP could also be obtained through the Path Collective Variables description coupled with metadynamics^43^. Alternatively to biased-MD simulation methods, certain spontaneous binding events can also be achieved through extensive standard MD simulations, for example, running for *μ*s to ms simulation time^44,45^. From MFEP we can extract information associated to the most probable trajectories in the CV space followed by the system involving between (meta-)stable states and it also allows us the determination of local and global transition states. In this way, we have traced the MFEP along the free energy landscape using the guidelines sketched in SI by means of the R-package metadynminer^46^. Several MFEP have been depicted in Figs. 6 and 7 where the global minimum is set to zero. The coordinates of minimum free energy paths are reported in Table 4 of SI.

Our results suggest that there exist many local and global stable states along on the complex 2D FEL reported in Figs. 6 and 7. For oncogenic KRas-4B-Far protein (Fig. 6), the most stable state (global minimum) and four more meta-stable states have been revealed and shown in Table 2.

**Table 2:** Minima on the FEL of oncogenic KRas-4B-Far (coordinates of CV given in nm). Each letter refers to one meta-stable state, with A standing for the global minimum of free energy when FAR is anchoring deeply into the membrane and GTP located near the interface. All locations of minima of FEL of Fig. 5 were obtained by using the metadynminer package.

| States | CV1 (gtp) | CV2 (far) | Free energy (kJ/mol) |
| --- | --- | --- | --- |
| A | -2.28 | -1.37 | 0.00 |
| B | -5.08 | -1.72 | 15.00 |
| C | -1.81 | -4.00 | 34.28 |
| D | -1.93 | -9.59 | 36.56 |
| E | -7.77 | -1.49 | 41.59 |

The stable or meta-stable configurations that oncogenic KRas-4B-Far may adopt when bound to the anionic membrane are as follows: in state A, FAR anchors into the “internal” regions of the membrane when GTP is close to the interface and binds the CD moiety as well; in state B, FAR anchors into the “internal” regions of membrane when GTP only binds the CD moiety; in state C FAR has been released off the membrane, but GTP is still bound at the interface; in state D, FAR is solvated by water molecules and GTP only binds the membrane interface and in state E, GTP reaches out further away from the lipidic region and is solvated by the aqueous solution, while FAR keeps being anchored into the membrane.

From 2D FEL we can also estimate the numerical values of the free energy barriers between the selected meta-stable configurations reported above:

1. While FAR is anchored to the membrane, GTP dissociates from stable state A then binds to the CD moiety (state B), requiring to cross a free energy barrier of 27.04 kJ/mol. The final state is essentially represented by configuration 3 of Fig. 5.
2. The difference between states C and D is most likely due to the location of HVR. HVR may bind the head groups of lipids (state C) and it can be solvated in the aqueous region (state E). The estimated free energy barrier ΔF between states C and D is of 17.20 kJ/mol.
3. We estimated a high free energy barrier of 42.44 kJ/mol from state A to state C. This explains that for oncogenic KRas-4B-Far, the affinity of FAR to the anionic membrane is extremely high.
4. ΔF for the transition between states B and E has been estimated to be of 27.5 kJ/mol, corresponding to the energy needed to get the release of GTP from its association from the CD of the protein to water solvation. This explains the high affinity of GTP to CD in the oncogenic KRas-4B-Far.

Correspondingly, in the case of the FEL of wild-type KRas-4B-Far protein, two global stable states (A, B) and two more meta-stable states (C, D) have been identified and shown in Table 3 whereas important MFEP are shown in Fig. 7.

**Table 3:** Minima on the FEL of wild-type KRas-4B-Far. Here each letter refers to one meta-stable state, among which A stands for the global state which is set to zero. Here locations of minima of FEL of Fig. 6 were obtained by using metadynminer package.

| States | CV1(gtp) | CV2(far) | Free energy (kJ/mol) |
| --- | --- | --- | --- |
| A | -4.06 | -3.49 | 0.00 |
| B | -3.94 | -1.93 | 1.92 |
| C | -3.35 | -0.97 | 5.14 |
| D | -3.82 | -5.91 | 15.25 |

In this case the main features are summarised as follows:

1. When FAR is bound to the membrane (state B), a barrier of 10.76 kJ/mol should be crossed by FAR shifting to the global stable state A while GTP keeps bound to the CD, but apart from the interface.
2. Shifting from state B to state C (FAR anchored in a deeper position and GTP closer to the interface) can be realised if a free energy barrier of 6.69 kJ/mol is surmounted.
3. When the system shifts its state between A and D (i.e. FAR solvated in the aqueous region), the crossing of a free energy barrier of 20.10 kJ/mol corresponds to the energy needed for the HVR to be released from the anionic membrane.
4. There are several meta-stable states when GTP locates ~2 nm away from the membrane center, indicating the existence of multiple configurations when GTP is around head groups of lipids, which may play a role in KRas-4B-Far signal transduction and interactions with other proteins *in vivo.*

### One-dimensional free energy profiles

From the two-dimensional well-tempered metadynamics simulation is possible to calculate one-dimensional (1D) free energy profiles. That kind of calculation allows us to directly compute free energy change along a single CV (*s_i_*, *i* = 1, 2), which can be directly compared to experimental findings^47^. 1D free energy profiles F(s_1_) for both systems (oncogenic, wild-type) after integrating out the second CV *s*_2_ have been represented in Fig. 8 and can be obtained according to Refs. ^47,48^ as follows:

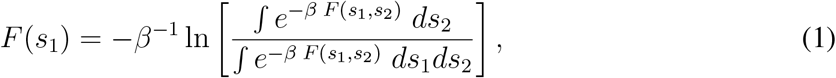

where *s*_1_ and *s*_2_ are the CV, *β* = 1/(*k*_B_*T*), *k*_B_ is the Boltzmann constant and T is the absolute temperature. This means that all possible paths for the CV labelled as *s*_2_ have been averaged. *F*(*s*_1_) reveals additional information about the most stable states only as a function of one CV.

From plots in Fig. 8 we can obtain meaningful free energy barriers related to particular processes related to movements of GTP and FAR along Z axis. The main findings are highlighted as follows:

1. In the case of the oncogenic KRas4B-Far system GTP needs to surmount a barrier of 30.53 kJ/mol to move from the interface of the bilayer to outside (A to B), in order to bind with the CD moiety.
2. The displacement of GTP from the interface of one leaflet of the membrane to the aqueous region requires about 49.49 kJ/mol (oncogenic) and 30.56 kJ/mol (wild-type).
3. In the same fashion, the free energy barrier to surmount for FAR release from the membrane (A) to be solvated in the aqueous region is of about 40 kJ/mol for the oncogenic case.
4. In its GTP-bound state, wild-type KRas-4B-Far could easily change its configurations between states A and B. In order to decline interactions between FAR and lipids, FAR crosses a barrier of 8.44 kJ/mol to reach state A from state B. The wild-type KRas-4B-Far interacts with the membrane firmly through its the HVR and FAR moieties. However, once its FAR dissociates from the membrane it has the ability to anchor back and act as a switch, which ensures its normal function. Accordingly, GTP is very stable in its location and can move from the interface to the CD moiety passing through barriers of less than 10 kJ/mol.
5. Comparing the free energy profiles of the oncogenic case with the ones for the wild-type protein, PHOS at Ser-181 strongly favors FAR anchoring to the membrane while GTP is located at the interface. This is consistent with our MD simulation results. This provides additional support to our statement that increasing its hydrophobic density through the carboxymethylation at Cys-185 does not always favor its protein recruitment to the PM when PHOS gets involved in the KRas-4B structure.

In a similar perspective, Zhou et al.^4^ published results of metadynamics simulations of KRas in plasma membranes on the conformational changes using very different collective variables: CV1 was the C*α*-atom RMS of residue 177-182 from a helical reference structure and CV2 the one end-to-end distance involving the C*α*-atom of residue 176 and 184, at the 1 *μ*s timescale. In such case, PHOS at Ser-181 of mutated KRas-G12V was found to favor a specific orientation state.

## 3 Discussion

KRas-4B belongs to a family of small GTPase that regulate cell growth, differentiation and survival which is frequently mutated in cancers such as in lung, colon and pancreatic ones. However the structural mechanisms at the atomic level of KRas-4B-membrane association are not well understood yet. Most efforts have been focused on the methylated KRas-4B-FMe protein after decades of research. However, a relatively high abundance of farnesylated KRas-4B-Far (wild-type and oncogenic) has been found in tumors. In this paper findings of three wild-type and mutated KRas-4B proteins obtained by using molecular dynamics and metadynamics simulations have been reported.

Firstly, we conducted MD simulations of three systems of GTP-bound KRas-4B (oncogenic KRas-4B-FMe and KRas-4B-Far, wild-type KRas-4B-Far) binding cell membranes constituted by DOPC (56%), DOPS (14%) and cholesterol (30%) including Cl solution at 310.15 K and at the fixed pressure of 1 atm. Three set of continuous MD simulations all reached the scale of 1000 ns. Each oncogenic KRas-4B protein contains two mutations: G12D and phosphorylation at its Ser-181 site. Our main interest was focused on the local structures and localisations of different moieties of KRas-4B proteins. As a general fact, KRas-4B association with the PM requires the penetration of FAR into the membrane and interactions of HVR with lipids in order to allow KRas-4B a proper localisation in the membrane. After a systematic analysis of meaningful data, we noted relevant differences between different cases. We observed that for wild-type KRas-4B-Far, demethylated Cys-185 allows spontaneous insertion and release of FAR into the anionic membrane bilayer without much difficulty. Conversely, oncogenic KRas-4B-Far keeps anchored to the membrane and staying in its active site in order to bind upstream regulators and downstream effectors. Further, oncogenic KRas-4B-FMe stays in an auto-inhibited state in which the HVR is sandwiched between the effector binding site of the CD and the membrane, blocking signal transduction pathways.

In order to characterise the system, microscopic properties such as the area per lipid, lipid thickness, radial distribution functions, penetration of FAR, GTP, etc. into the membrane and deuterium order parameters have been evaluated. Interactions, such as HB and long-lived salt bridges between different active sites of GTP, the CD, the HVR and lipids have been revealed that to play a central role in the stabilisation of KRas-4B membrane proteins. In particular, only in the oncogenic KRas-4B-Far case long-lived salt bridges have been located between oxygen and hydrogen atoms from FAR and PHOS, providing large stability to the anchoring of the KRas-4B-Far into the PM. We believe this is the key for the permanent association between the oncogenic protein and the cell membrane, which will happen in the inner leaflet of real membranes. Further, the information reported in this work could be very valuable to the development of farnesyltransferase inhibitors as anti-cancer agents^49^.

Among the two chosen mutations to generate the oncogenic KRas-4B proteins, we have observed that PHOS has important influence on the behavior of oncogenic KRas-4B proteins, together with the effect of PTM such as carboxymethylation, which always play a role in the localisation of KRas-4B at the PM. By investigating the penetration along the membrane normal of different moieties of the KRas-4B structures, we have revealed that bigger distances of the CD to the membrane center help to expose specific sites of CD to bind upstream regulators and downstream effectors, finally populating their active state for the mutated proteins.

To the best of our knowledge, the free energy profiles of GTP binding to anionic membrane bilayers have been reported for the first time, giving first-hand information about FEL of GTP-bound wild-type/oncogenic KRas-4B-Far binding to anionic membrane systems. With the help of well-tempered metadynamics simulations we have calculated 2D free energy landscapes and the 1D free energy profiles along one CV. We have located global and local stable states of the two systems that have been chosen to elucidate the joint effect of PHOS and G12D on the KRas-4B-membrane binding. Two CV have been taken into consideration to describe the conformational changes along free energy paths. CV1 is the distance between the center of mass of GTP and the center of the membrane, and CV2 is the distance between the center of mass of farnesyl group of Cys-185 and the center of the membrane along the normal direction.

Our results indicate that for GTP-bound oncogenic KRas-4B-Far it exists a free energy barrier of 42.44 kJ/mol for the departure of FAR from the PM, which explains the large stability of the FAR moiety of oncogenic KRas-4B-Far at the anionic membrane. This corresponds to the configuration 3 of Fig. 5, characteristical from MD simulations. Correspondingly, a barrier of 10.76 kJ/mol is likely to be crossed for FAR of wild-type KRas-4B-Far shifting from anchoring to the PM to being solvated by water while GTP keeps bound to the CD (shift between configurations 1 and 2 of Fig. 5). We have estimated a free energy barrier of 20.10 kJ/mol when FAR of the wild-type case shifts from ~4 nm to 6 nm away from the membrane center, corresponding to the energies needed for the HVR dissociating from the anionic membrane.

From the calculation of 1D integrated free energy profiles of both systems, important observations have been presented. GTP needs 30.53 kJ/mol to move from the interface of the bilayer to bind with the CD moiety for oncogenic KRas-4B-Far system, whereas only 7.74 kJ/mol free energy are required for the wild-type case. Similar barriers of 33.98 kJ/mol and 35.36 kJ/mol for oncogenic and wild-type system, respectively, need to be surmounted for GTP to dissociate from the CD to be solvated by water molecules, indicating that mutation at G12D does not have significant influence on the dynamics of GTP. Most importantly, the free energy needed for FAR release from the membrane to be solvated in the aqueous region is of 46.01 kJ/mol for oncogenic case and of 15.16 kJ/mol for wild-type case, respectively. This provides additional support to the crucial difference between the two species directly related with the strong bonding between FAR and the PHOS group.

## 4 Methods

We conducted MD simulations of model anionic (neutral) cell membranes constituted by DOPC (56%), DOPS (14%) and cholesterol (30%) for the three KRas-4B isoforms with sequences represented in Fig. 1 of SI. Each system contains a total of 304 lipid molecules fully solvated by 60,000 TIP3P water molecules^50^ in potassium chloride solution at the human body concentration (0.15 M), yielding a system size of 222,000 atoms. All MD inputs were generated using CHARMM-GUI membrane builder^51,52^ and the CHARMM36m force field^53^ was adopted for lipid-lipid and lipid-protein interactions. Crystal structure of KRas-4B with partially disordered hypervariable region (pdb 5TB5) and GTP (pdb 5VQ2) were used to generate full length GTP-bound KRas-4B proteins. Both pdb files were downloaded from RCSB PDB Protein Data Bank^54^. The three sets of full length KRas-4B and GTP were solvated in a water box and equilibrated for 20 ns before generating the full setups. KRas-4B was initially set anchoring to membrane in the beginning of simulation in each system. All systems were energy minimised for 5000 steps followed by three 250 ps simulations and then four additional 500 ps equilibrium runs while gradually reducing the harmonic constraints on the systems. Production runs were performed with an NPT ensemble for 1 *μ*s. The pressure and temperature were set at 1 atm and 310.15 K respectively, well above the corresponding transition temperature for DOPC and DOPS, in order to ensure that we were simulating the liquid crystalline state. In all MD simulations, the GROMACS/2018.3 package was employed^55^. Time steps of 2 fs were used in all production simulations and the particle mesh Ewald method with Coulomb radius of 1.2 nm was employed to compute long ranged electrostatic interactions. The cutoff for Lennard-Jones interactions was set to 1.2 nm. Pressure was controlled by a Parrinello-Rahman piston with damping coefficient of 5 ps^−1^ whereas temperature was controlled by a Nosé-Hoover thermostat with a damping coefficient of 1 ps^−1^. Periodic boundary conditions in three directions of space have been taken.

After equilibrating properly the system in the two cases of wild-type/oncogenic KRas-4B-Far from a 1 μs run for each system, we switched to run another 1.1 *μ*s of well-tempered metadynamics simulations to perform Gibbs free energy calculations of KRas-4B binding at anionic phospholipid membrane bilayers, starting from the last configuration of MD simulations. Undoubtedly, the problem of computing free energy landscapes in multidimensional systems has been extensively discussed in the literature. Among the wide variety of methods proposed, we can highlight transition path sampling^56–60^, adaptive biasing force^61^ or umbrella sampling methods^62,63^, to mention a few. In the present work we have employed well-tempered metadynamics, a method able to efficiently explore free energy surfaces of complex systems using multiple reaction coordinates in a very successful way for a wide variety of complex systems^64–68^.

Well-tempered metadynamics simulations were performed using the joint GROMACS/2018.3-plumed tool^69^. The isothermal-isobaric NPT ensemble at temperature 310.15 K and pressure 1 atm was adopted in all cases. Periodic boundary conditions in the three directions of space were also considered. Two collective variables were considered to describe the conformational changes along free energy paths. CV1 is the distance ‘gtp’ between the center of mass of GTP and the center of the membrane (i.e. *z* = 0) and CV2 is the distance ‘far’ between the center of mass of the farnesyl group of Cys-185 and the center of the membrane, where “wt.” and “onc.” represent the “wild-type” and “oncogenic” classes, respectively. Parameters of well-tempered metadynamics simulations are listed in Table 3 of SI.

## Acknowledgements

We warmly thank Profs. Carles Calero, Giancarlo Franzese and Dr. Martin Goethe for fruitful discussions. We also thank financial support provided by the Spanish Ministry of Science, Innovation and Universities (project number PGC2018-099277-B-C21, funds MCIU/AEI/FEDER, UE). Huixia Lu is a Ph.D. fellow from the Chinese Scholarship Council (grant 201607040059). Computational resources awarded by the Barcelona Supercomputing Center-Spanish Supercomputing Network (grants FI-2018-3-0023, FI-2019-1-0004, FI-2019-2-0004 and FI-2019-3-0008) are also acknowledged.

