## Supplementary Information for "Long-lasting salt bridges provide the anchoring mechanism of oncogenic KRas-4B proteins at cell membranes"

**Huixia Lu**[1,+] **and Jordi Martí**[1,*,+]

[1]Department of Physics, Technical University of Catalonia-Barcelona Tech, B4-B5 UPC Northern Campus, Barcelona, Catalonia, Spain
[*]
[+]these authors contributed equally to this work

### ABSTRACT

Here we report full computational details on: (1) system structure and composition; (2) molecular dynamics simulation setups for the full KRas-4B/DOPC/DOPS/cholesterol membrane system; (3) supplementary results on structure of the membrane as well as radial distribution functions related to the catalytic domain and hypervariable regions of the protein; (4) well-tempered metadynamics simulations that lead to the construction of the 2D free energy surfaces as a function of the class of protein, focusing especially on their stability and convergence; (5) technical details about the calculation of minimum free energy paths.

### 1 Supplementary Notes

#### 1.1 Initial setups and data for molecular dynamics simulations

The three classes of KRas-4B proteins that have been considered in this work are reported in Fig. 1 (right). Their component aminoacids are listed in Table 1 and the sketches of parts of the full systems are described in Fig. 2. The process of post-translation modifications (PTM) required to allow KRas-4B to function properly in cells is represented in Fig. 1 (left): Firstly the prenylation reaction, catalysed by cytosolic farnesyltrasferase (FTase) or geranylgeranyltransferase (GGTase), proceeds through the addition of an isoprenyl group to the Cys-185 side chain. Then farnesylated KRas-4B is ready for further processing: hydrolysis, catalysed by the endopeptidase enzyme called Ras-converting enzyme 1 (RCE1), during the process the VIM motif (HVR tail composed of three amino-acids: valine-isoleucine-methionine) of the C-terminal Cys-185 is lost in step 2. Later KRas-4B is transferred to the endoplasmic reticulum for carboxymethylation at the carboxyl terminus of Cys-185 catalysed by isoprenylcysteine carboxyl methyltransferase (ICMT), forming a reversible ester bond. The outcome of these modifications is the farnesylated and methylated KRas-4B (KRas-4B-FMe). The reversible ester bond can go through decarboxymethylation, catalysed by prenylated/polyisoprenylated methylated protein methyl esterases (PMPEases) giving rise to a farnesylated and demethylated KRas-4B (KRas-4B-Far) which is the product of step 2 and reactant of step 3. This reversible reaction can modulate the equilibrium of methylated/demethylated KRas-4B population in tumors and consequently can impact downstream signalling, protein-protein interactions, or protein-lipid interactions[1].

A deuterium order parameter $S_{CD}$ was defined[2–5] for each $CH_2$ and CH groups of the DOPC lipid tails as Eq. 1 and its averaged results are shown in Fig. 3 for both tail chains of all DOPC lipids in all KRas-4B systems studied in this work.

$$S_{CD} = \frac{1}{2}(3 < \cos^2\theta_{CH} > -1), \tag{1}$$

where $\theta_{CH}$ is the angle between the membrane normal and a CH-bond. Brackets in Eq.1 indicate ensemble average for all lipids. $S_{CD}$ can be also obtained from $^2$H NMR experiments[6].

We have considered radial distribution functions (RDF) between highlighted sites of molecules (see Fig. 2) and involved sites of the catalytic domain and the hypervariable region of KRas, together with the corresponding molecular structure of aminoacids of KRas-4B (see Figs. 4 and 5). The RDF are presented in Figs. 6 and 7 for CD and HVR sites, respectively.

The distribution of distances between oxygen and hydrogens of FAR and corresponding sites in the phosphate group of the PHOS is displayed in Fig. 8. The localisation of FAR, GTP, CD and HVR referenced to the positions of the interfaces of the membranes are represented in Fig. 9.

### 1.2 Convergence of the well-tempered metadynamics simulations

Technical details of the metadynamics simulations were described in Section 4 of the main text. The values for the parameters of the well-tempered metadynamics simulations[7] are listed in Table 3.

First of all, convergence of well-tempered metadynamics simulations depends on the number of transition events between states and sampling of all physically interesting regions of the collective variables (CV)[8]. The challenge of well-tempered metadynamics is well known, especially when applied to protein-bilayers systems with hundreds of thousands of atoms, in order to sample all relevant conformations and the full range of CV. Our work is based on the two selected CV defined in Section 4 of main text (CV1 is the distance between the center of mass of GTP and z = 0, and CV2 is the distance between the center of mass of the farnesyl group of Cys-185 and z = 0, where Z-axis is defined along the direction normal to the membrane).

It is usual to monitor the size of the hills of the Gaussian kernels deposited along the simulation. As the simulation progresses and the added bias grows, the Gaussian height is progressively reduced, eventually including low-height spikes. We can visualise the decrease of the Gaussian height during the simulation. In the long term, the Gaussian height becomes smaller and smaller while the system diffuses in the entire CV space. The height of the biased potential decreased accordingly along the simulation runs, as indicated in Fig. 10. In all cases, a quasi-flat profile is already seen after 600 ns, although some high spikes are appearing due to large fluctuations in the values of CV when covering all configurational space.

Nevertheless, the fact that the Gaussian height is decreasing to zero should not be used as a unique measure of convergence of a metadynamics simulation. By inspecting Fig. 11 where the well-tempered metadynamics trajectories of the two CV (gtp and far) are reported along the full time span of our simulations, we can see that the two systems were both initialised in one of their (meta-)stable states. After $\sim 0.1$ ns, the systems were pushed by the metadynamics bias potential to visit other local minima. As simulations continue, the bias potential fills the underlying free energy landscape, and both CV are able to diffuse in full phase space along the final 1050 ns simulation time span.

We can clearly identify that for both systems GTP has diffused efficiently in the full collective variable space except for the center of the membrane. No permeation of GTP across the membrane has been observed, which gives an indication that the corresponding free energy barrier is too high for GTP. In the case of the oncogenic KRas-4B-Far system, FAR visited all the possible space in one leaflet. However, for wild-type KRas-4B-Far system, FAR only reached out in the CV space of around 7 nm away from the membrane center along the membrane normal direction, which is enough for our aim of exploring FAR release from the membrane and describing its FEL covering all the relevant CV space.

Convergence can be evaluated in different ways, such as monitoring the stability of free energy barriers between different states or by plotting integrated free energy profiles along the simulation time. We have analysed convergence by two ways: (1) reporting the time cumulative average of 1D free energy profiles along the simulation time, and (2) calculating the free-energy difference between the two local minima in the one-dimensional free energy along each CV as a function of simulation time. At convergence, the reconstructed free energy profiles should be similar, apart from a constant offset. From the results of Fig. 12, we can see that 1D free energy profiles are very similar after simulation time of 800 ns for CV 'gtp' and 900 ns for CV 'far' in the oncogenic system. For the wild-type system, convergence was realised much earlier, since 600 ns for gtp and 700 ns for far.

To assess the convergence of the simulation more quantitatively, we have also calculated the evolution of free energy barriers ($\Delta F$) between two selected free energy basins representing two meta-stable configurations for KRas-4B-Far as a function of simulation time, whose results are reported in Fig. 13. There we can observe that $\Delta F$ between the two chosen free energy basins tends to be similar after long cumulative time spans which leads us to fully converged free energies for the two CV considered in this work. The estimated free energy profiles and corresponding $\Delta F$s do not change significantly in the final part of our simulations which also ensures us that the simulations were fully converged.

### 1.3 Calculation of the minimum free energy paths

In order to obtain and evaluate the paths connecting two meta-stable states located on the 2D free energy surfaces reported in the Section "Two-dimensional free energy landscapes", we have considered a process including three steps: (a) fixing two local minima; (b) locating a coarse path and (c) refining the path. In all calculations reported in the present work we reported 8 and 10 points per path. The results for a series of paths between consecutive minima have been included in Figures 5 and 6 of the manuscript and their coordinates are numerically reported in Table 4. The first coordinate of each point corresponds to CV1 and the final coordinate to CV2. The R-package metadynminer[9] reads HILLS files from PLUMED, calculates free energy surface by fast Bias Sum algorithm, finds minima and analyses transition paths by Nudged Elastic Band method.

### 2 Supplementary Tables

**Table 1.** Aminoacid components of the KRas-4B proteins. Abbreviations as used in the text and in several figures.

| Full name | Abbreviation |
|---|---|
| Alanine | Ala |
| Arginine | Arg |
| Asparagine | Asn |
| Aspartate | Asp |
| Cysteine | Cys |
| Glutamate | Glu |
| Glutamine | Gln |
| Glycine | Gly |
| Histidine | His |
| Isoleucine | Ile |
| Leucine | Leu |
| Lysine | Lys |
| Methionine | Met |
| Phenylalanine | Phe |
| Proline | Pro |
| Serine | Ser |
| Threonine | Thr |
| Tryptophan | Trp |
| Tyrosine | Tyr |
| Valine | Val |

**Table 2.** Area per lipid ($A$) and thickness ($\Delta z$) of the anionic membrane for all the KRas-4B systems studied in this work. Estimated errors in parenthesis.

| System | $A$ ($nm^2$) | $\Delta z$ (nm) |
|---|---|---|
| wt. KRas-4B-Far | 0.523 (0.007) | 4.35 (0.05) |
| onc. KRas-4B-Far | 0.525 (0.006) | 4.23 (0.04) |
| onc. KRas-4B-FMe | 0.524 (0.006) | 4.34 (0.05) |

**Table 3.** Parameters employed in metadynamics simulations.

| System | onc. KRas-4B-Far | wt. KRas-4B-Far |
|---|---|---|
| Gaussian width of CV1 [nm] | 0.10 | 0.10 |
| Gaussian width of CV2 [nm] | 0.35 | 0.35 |
| Starting (Gaussian) hill [kJ/mol] | 2.0 | 1.2 |
| Deposition stride [ps] | 1 | 1 |
| Bias factor | 10 | 5 |
| Simulation time [ns] | 1100 | 1050 |

**Table 4.** Coordinates of segments forming selected paths for oncogenic and wild-type KRas-4B-Far systems. Along each path 8 locations of free energy spots (fespot, in kJ/mol) are displayed.

| Minimum free energy paths | | | | | | | |
|---|---|---|---|---|---|---|---|
| onc. KRas-4B-Far | | | | wt. KRas-4B-Far | | | |
| Stable states | path x | path y | fespot | Stable states | path x | path y | fespot |
| B | -5.08 | -1.72 | 15.00 | C | -3.35 | -0.97 | 5.14 |
| | -4.82 | -1.24 | 19.50 | | -3.36 | -1.07 | 7.00 |
| | -4.03 | -1.34 | 27.04 | | -3.80 | -1.10 | 8.61 |
| | -2.68 | -1.67 | 11.55 | | -3.79 | -1.18 | 8.02 |
| | -2.29 | -1.55 | 1.39 | | -3.77 | -1.25 | 7.68 |
| | -2.00 | -1.59 | 1.79 | | -3.76 | -1.41 | 6.69 |
| | -1.89 | -1.64 | 0.36 | | -3.76 | -1.63 | 4.06 |
| | -2.01 | -1.61 | 1.78 | | -3.78 | -1.77 | 2.48 |
| | -2.19 | -1.49 | 0.65 | | -3.75 | -1.88 | 1.92 |
| A | -2.28 | -1.37 | 0.00 | B | -3.94 | -1.93 | 1.92 |
| | -2.05 | -1.44 | 2.48 | | -3.80 | -1.93 | 2.02 |
| | -2.07 | -1.69 | 2.25 | | -3.77 | -1.99 | 2.53 |
| | -1.92 | -1.61 | 0.36 | | -3.95 | -2.27 | 9.43 |
| | -1.98 | -1.78 | 4.48 | | -3.64 | -2.61 | 10.76 |
| | -1.98 | -1.97 | 8.38 | | -4.05 | -3.00 | 8.29 |
| | -1.99 | -2.34 | 24.97 | | -3.94 | -3.48 | 0.04 |
| | -1.95 | -2.86 | 35.40 | | -3.96 | -3.50 | 0.04 |
| | -1.63 | -3.33 | 42.44 | | -3.93 | -3.49 | 0.04 |
| C | -1.81 | -4.00 | 34.28 | A | -4.06 | -3.49 | 0.00 |
| | -1.75 | -3.93 | 34.28 | | -3.71 | -3.75 | 6.27 |
| | -1.62 | -4.04 | 41.80 | | -3.92 | -3.98 | 7.10 |
| | -2.28 | -5.03 | 47.22 | | -4.07 | -4.26 | 10.73 |
| | -2.11 | -5.36 | 44.82 | | -3.92 | -4.78 | 13.13 |
| | -1.98 | -6.43 | 40.92 | | -3.88 | -5.04 | 15.57 |
| | -2.06 | -7.54 | 51.48 | | -3.73 | -5.30 | 16.93 |
| | -2.38 | -8.51 | 45.08 | | -3.75 | -5.51 | 20.10 |
| | -2.05 | -9.13 | 42.67 | | -3.76 | -5.67 | 20.04 |
| D | -1.93 | -9.59 | 36.56 | D | -3.82 | -5.91 | 15.25 |

### 3 Supplementary Figures

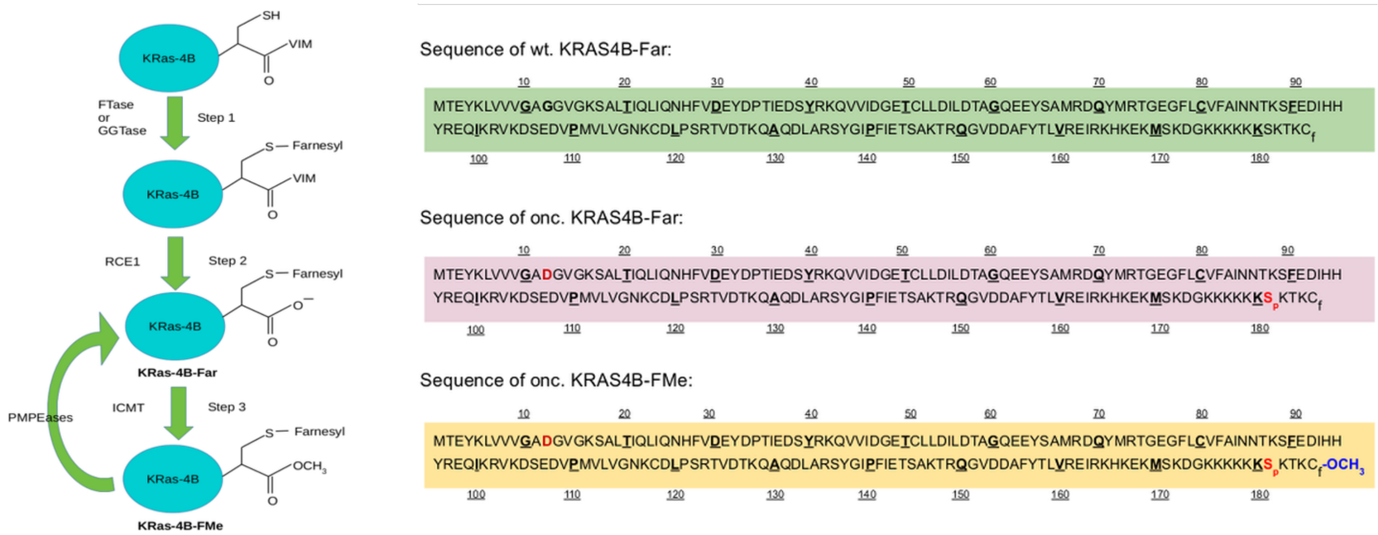


**Figure 1.** Process of post-translation modifications responsible to the formation of KRas-4B-Far and KRas-4B-FMe (left). At the right side, three sequences of different KRas-4B structures. Mutated sites are in red color. Here $C_f$ denotes the farnesylated Cys-185 site and $S_p$ represents the phosphorylated Ser-181 site. Methyl group of KRas-4B-FMe is in blue color.

**9.** Hošek, P. & Spiwok, V. Metadyn view: Fast web-based viewer of free energy surfaces calculated by metadynamics. *Comput. Phys. Commun.* **198**, 222–229 (2016).

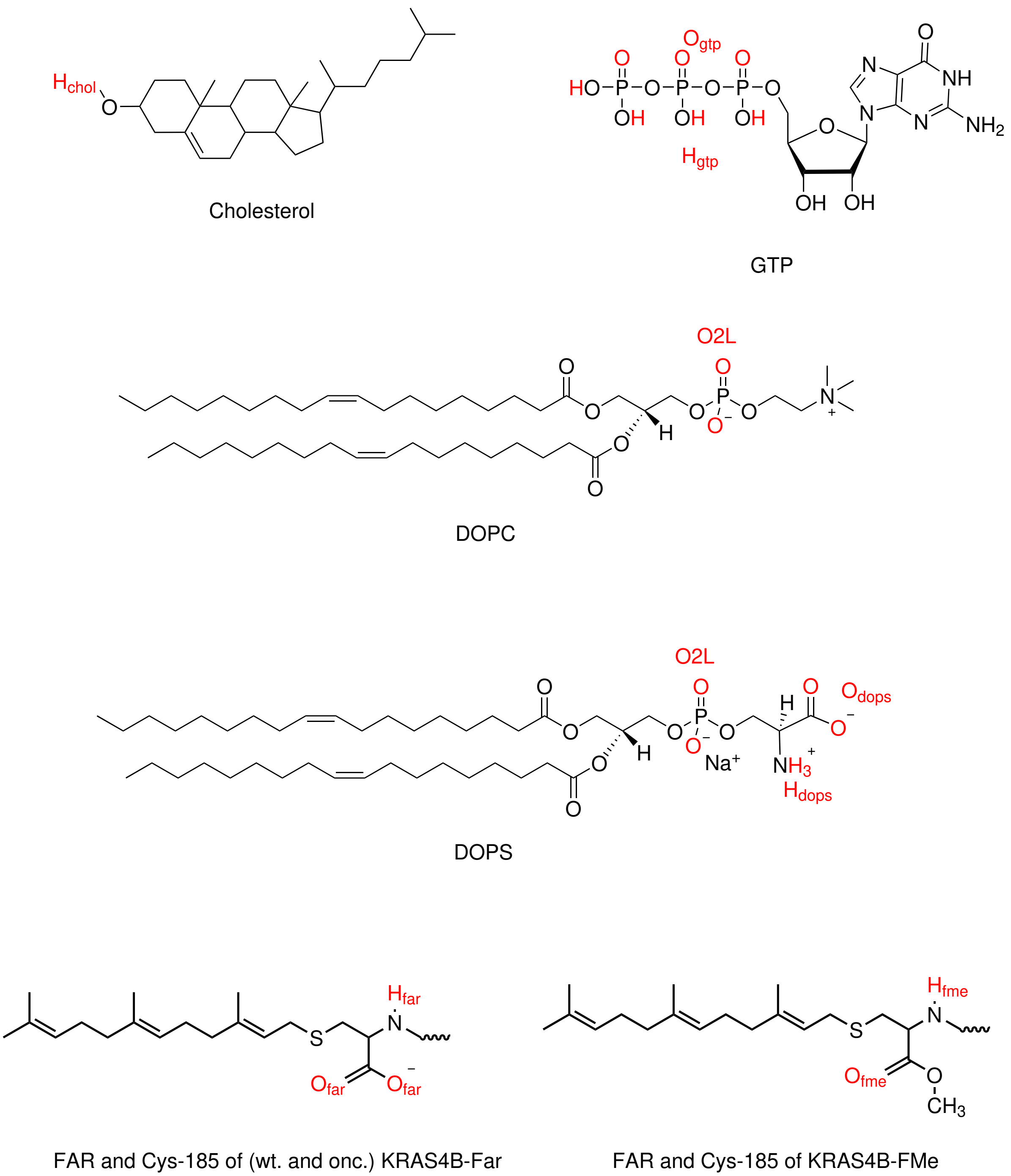


**Figure 2.** Sketches of GTP, DOPC, DOPS, cholesterol, and farnesyl in its phosphorylated and methylated forms.

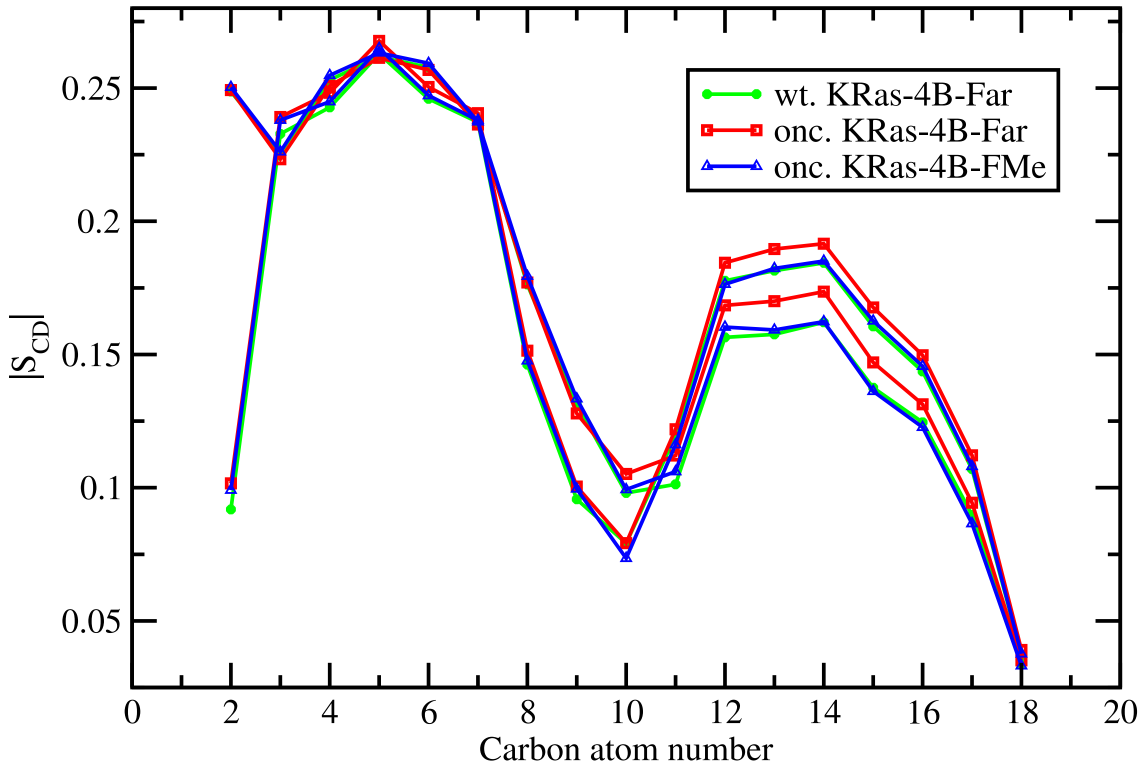


**Figure 3.** Averaged $S_{CD}$ for sn-1, sn-2 chains of DOPC in all systems.

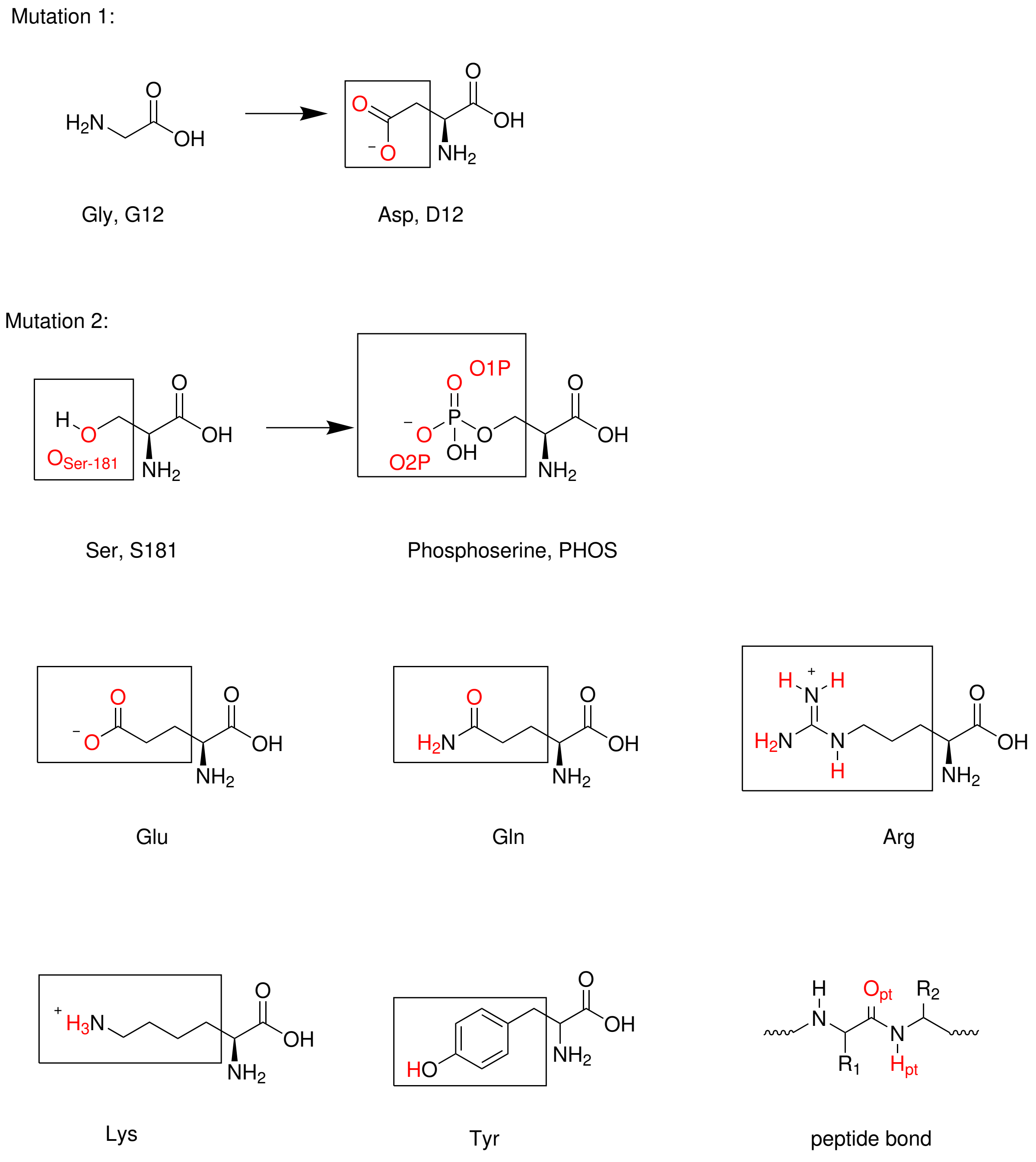


**Figure 4.** Two mutated sites in structures of onc. KRas-4B-Far/KRas-4B-FMe and active aminoacids of KRas-4B with their side chains framed.

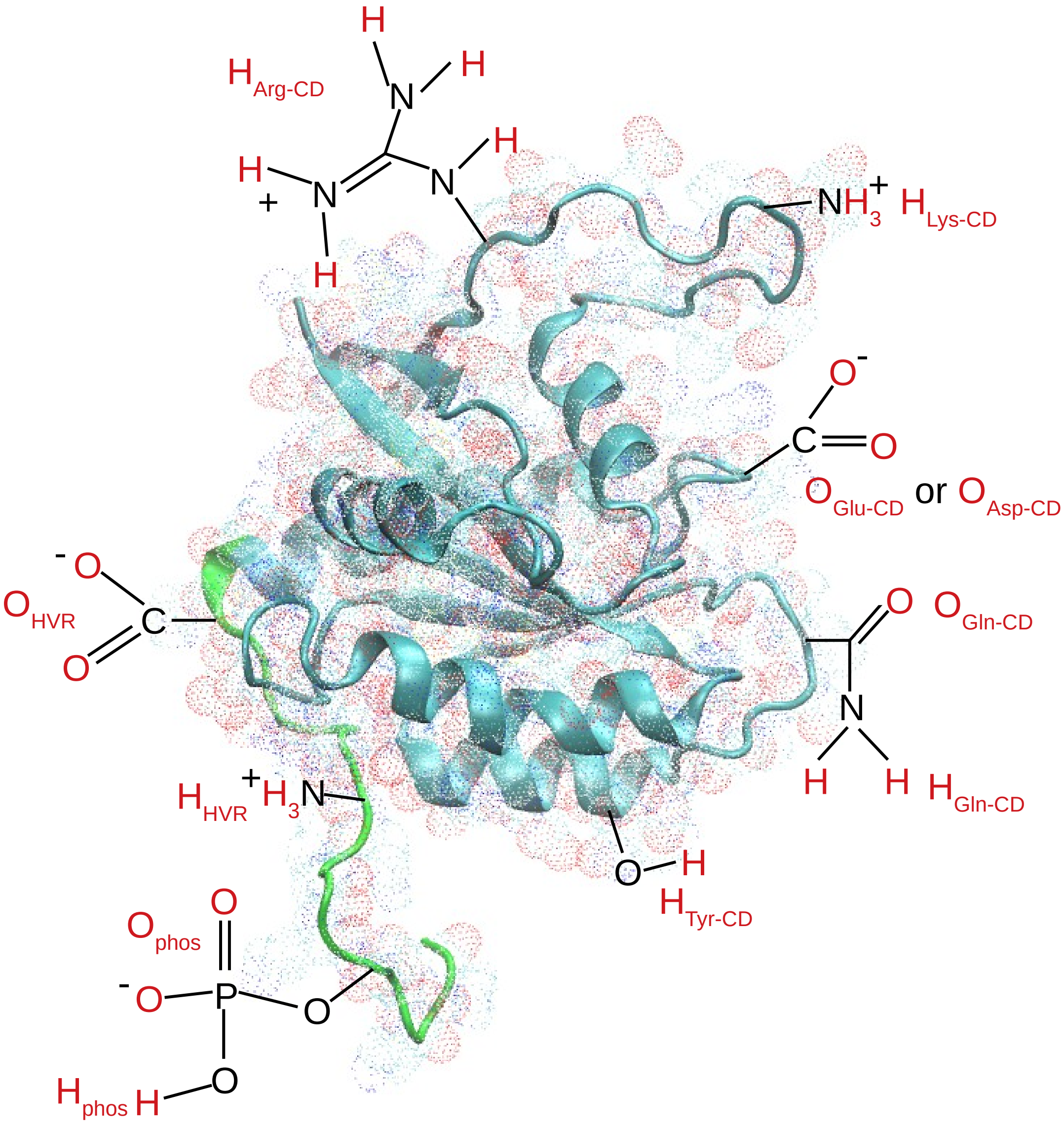


**Figure 5.** Selected active sites of the CD (cyan) and the HVR (green) of KRas-4B are depicted in red color. Farnesyl not shown. Thereinto, $H_{Arg-CD}$ donates the cationic hydrogen atoms from the guanidinium ($RNHC(NH_2)_2^+$) of arginine of the CD that share the same atom type, $H_{Lys-CD}$ and $H_{HVR}$ represent the cationic ammonium ($RNH_3{}^+$) from lysine of the CD and of the HVR. $O_{Glu-CD}$ and $O_{Asp-CD}$ represent oxygen atoms of the anionic carboxylate ($RCOO^-$) of glutamate aminoacid and aspartate aminoacids of the CD, respectively. $O_{HVR}$ represents all oxygen atoms of the anionic carboxylate ($RCOO^-$) of the HVR. $O_{Gln-CD}$ and $O_{Gln-CD}$ donate active oxygen and hydrogen atoms of side chain of glutamine aminoacid of the CD. Moreover, $O_{phos}$ and $H_{phos}$ donate two active oxygen atoms and one hydrogen atom of the phosphate group from PHOS.

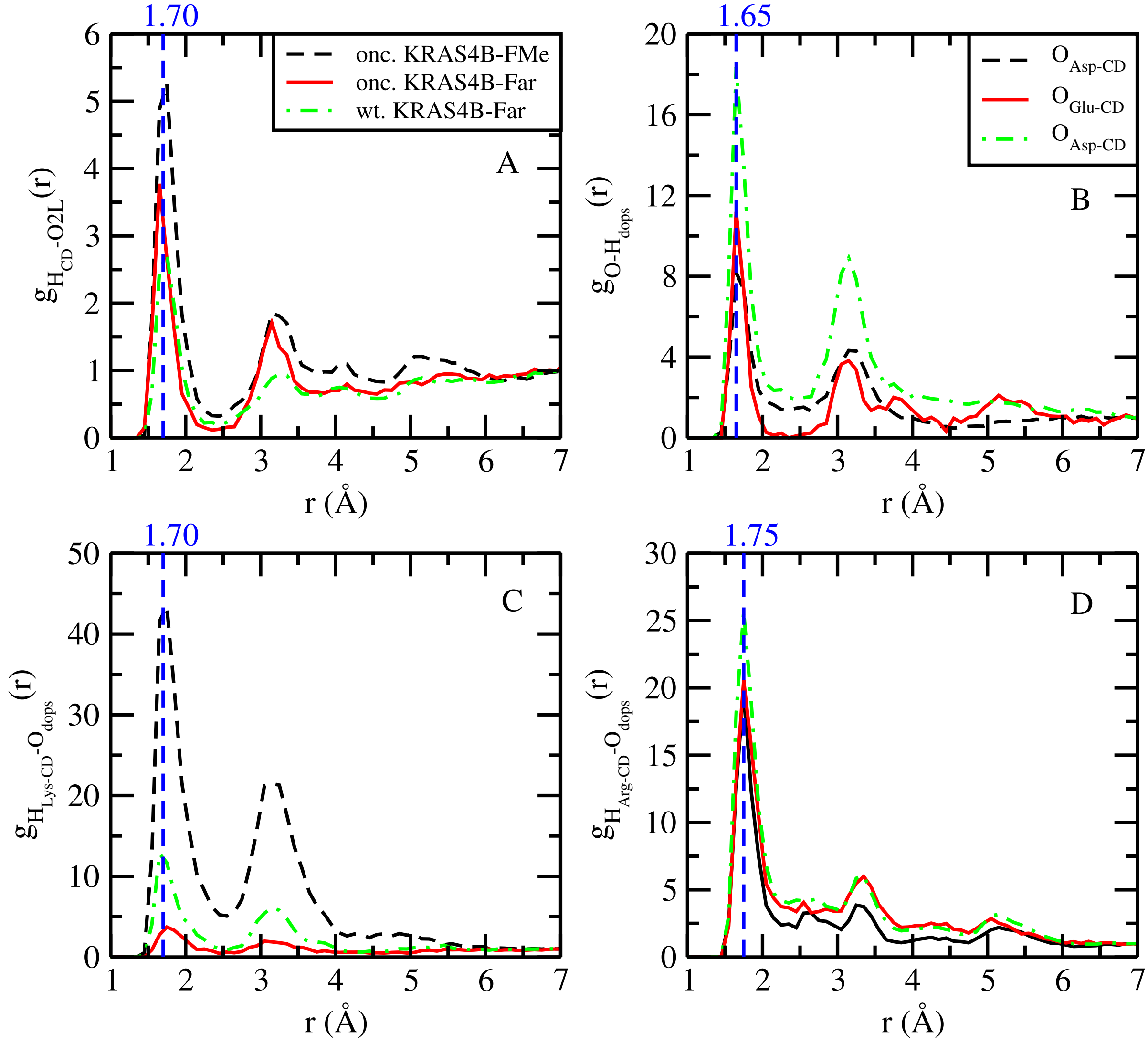


**Figure 6.** Selected RDF for selected sites of the CD with head groups of lipids. In all panels of this figure, as indicated in panel A, we use three colors to represent three systems in order to avoid the duplication and repetition, same as Fig. 7. Thereinto, $H_{CD}$ from panel A refers to the averaged value of $H_{Arg-CD}$ and $H_{Lys-CD}$.

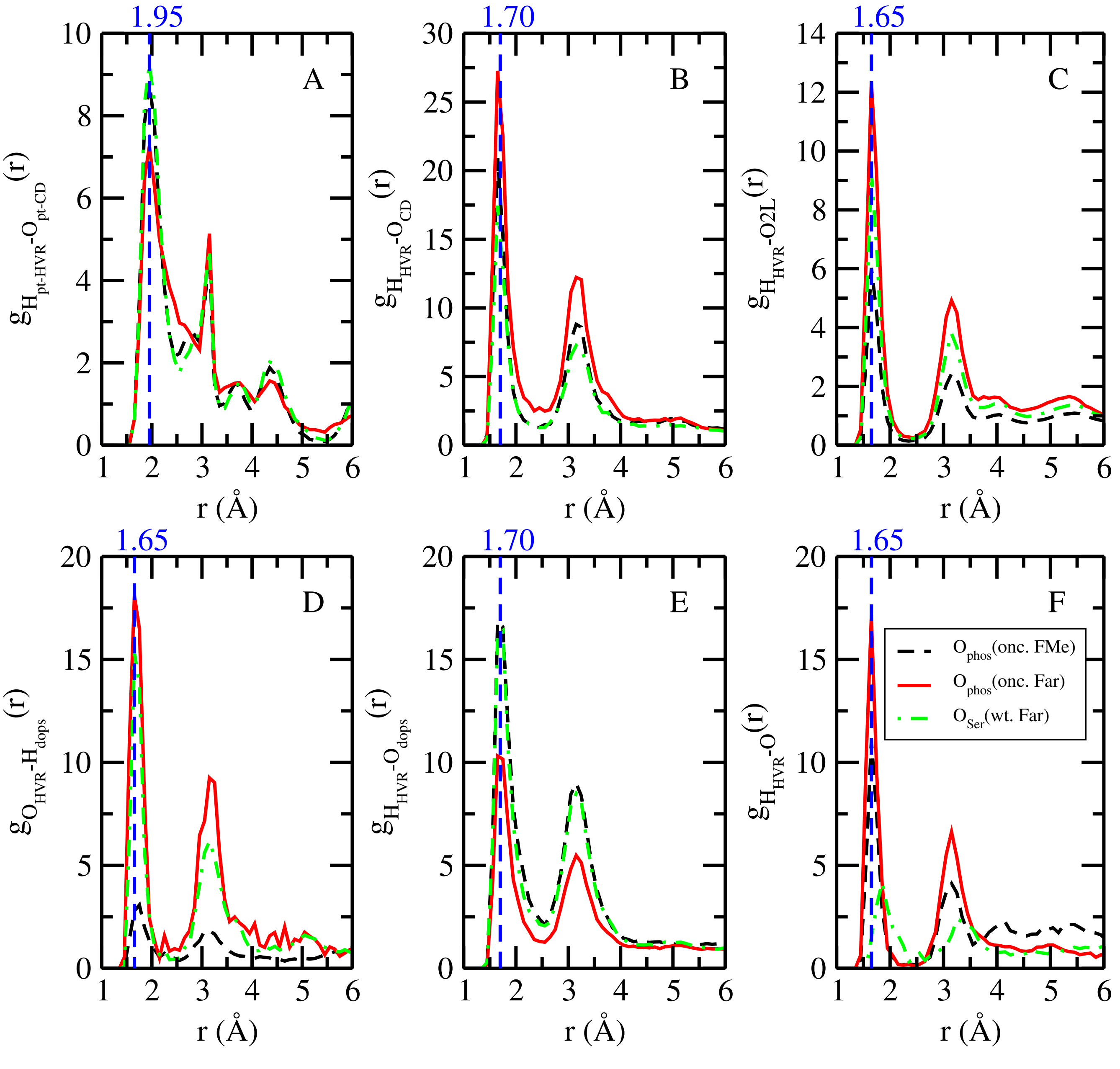


**Figure 7.** Selected RDF for selected sites of the HVR with atoms of lipids and the CD. Here $O_{pt-CD}$ and $H_{pt-HVR}$ represent oxygen and hydrogen atoms of the peptide bonds from the CD and the HVR. And $O_{CD}$ stands for oxygen atoms of $O_{Glu-CD}$ and $O_{Asp-CD}$ from the CD.

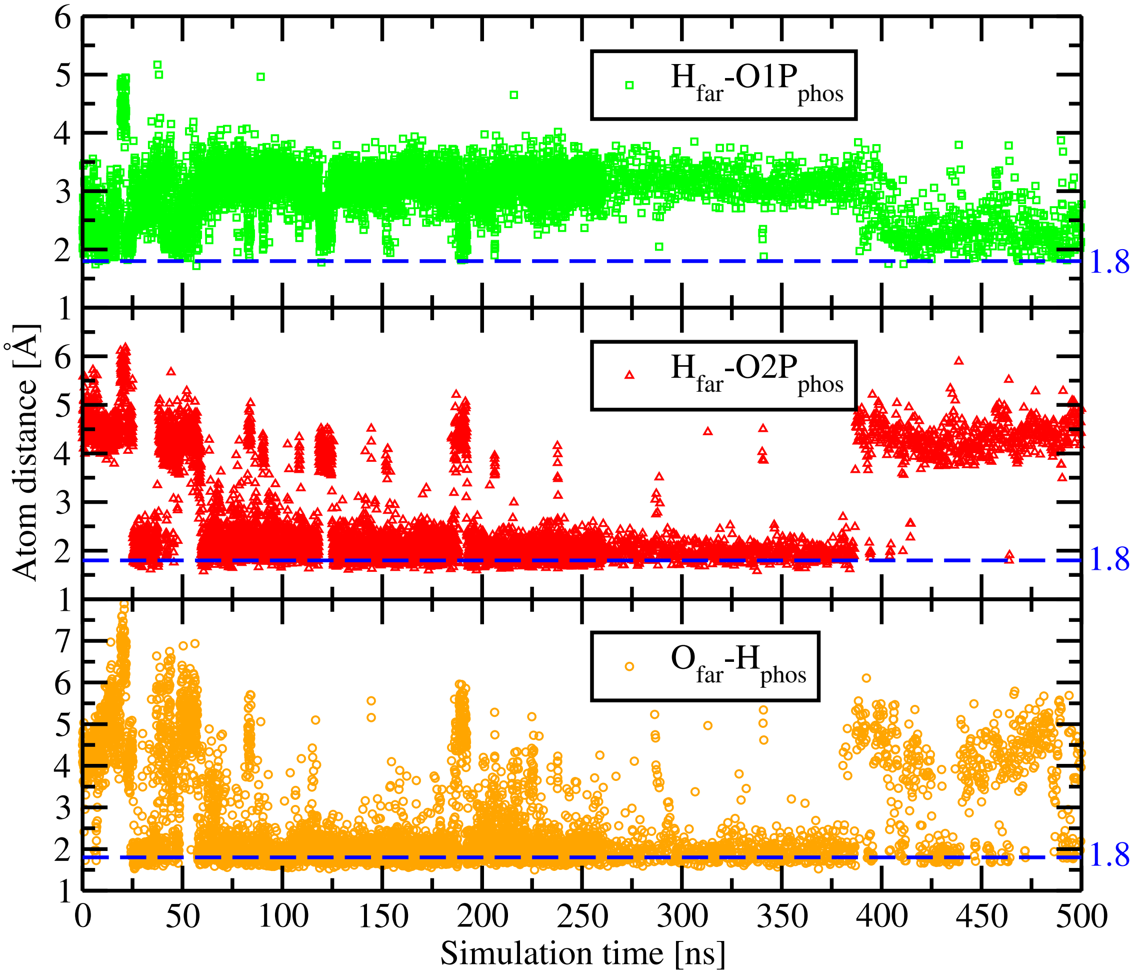


**Figure 8.** Distance distribution of selected sites of FAR and PHOS as a function of simulation time (see Fig. 3 of main text). Blue dashed lines indicate the typical HB distance (1.8 Å). O1P and O2P represent the two oxygen atoms belong to phosphate group of PHOS (see Fig. 4).

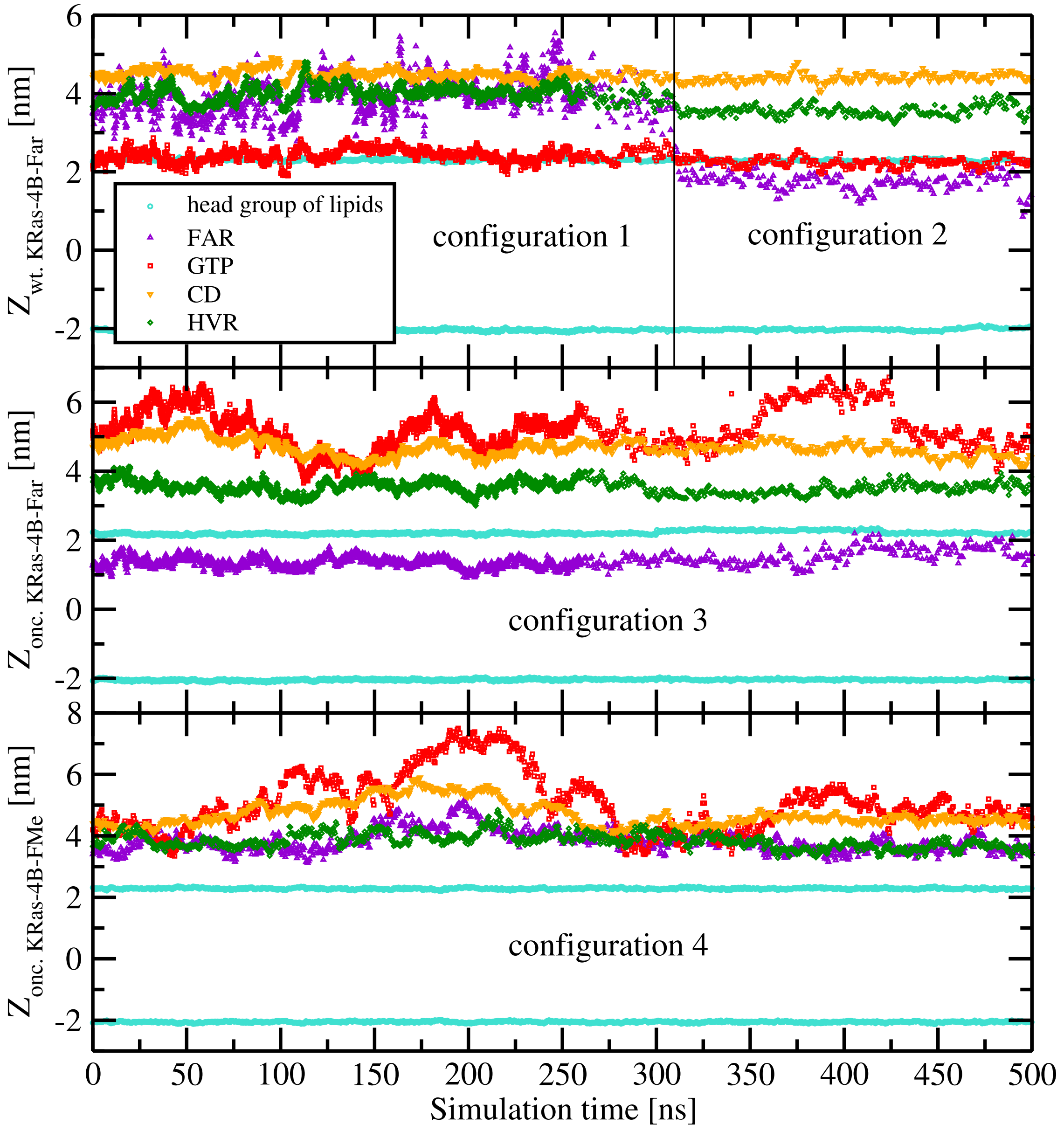


**Figure 9.** Localisation of KRas-4B domains and GTP as a function of the membrane normal coordinate *versus* simulation time. Geometric centers of the catalytic domain (triangles down, orange), the HVR (diamonds, green), FAR (triangles up, violet), GTP (squares, red) and phosphorus atoms of DOPC lipids from both leaflets (cyan, circles). The four favored configurations described here are reported in Fig. 5 of the main text.

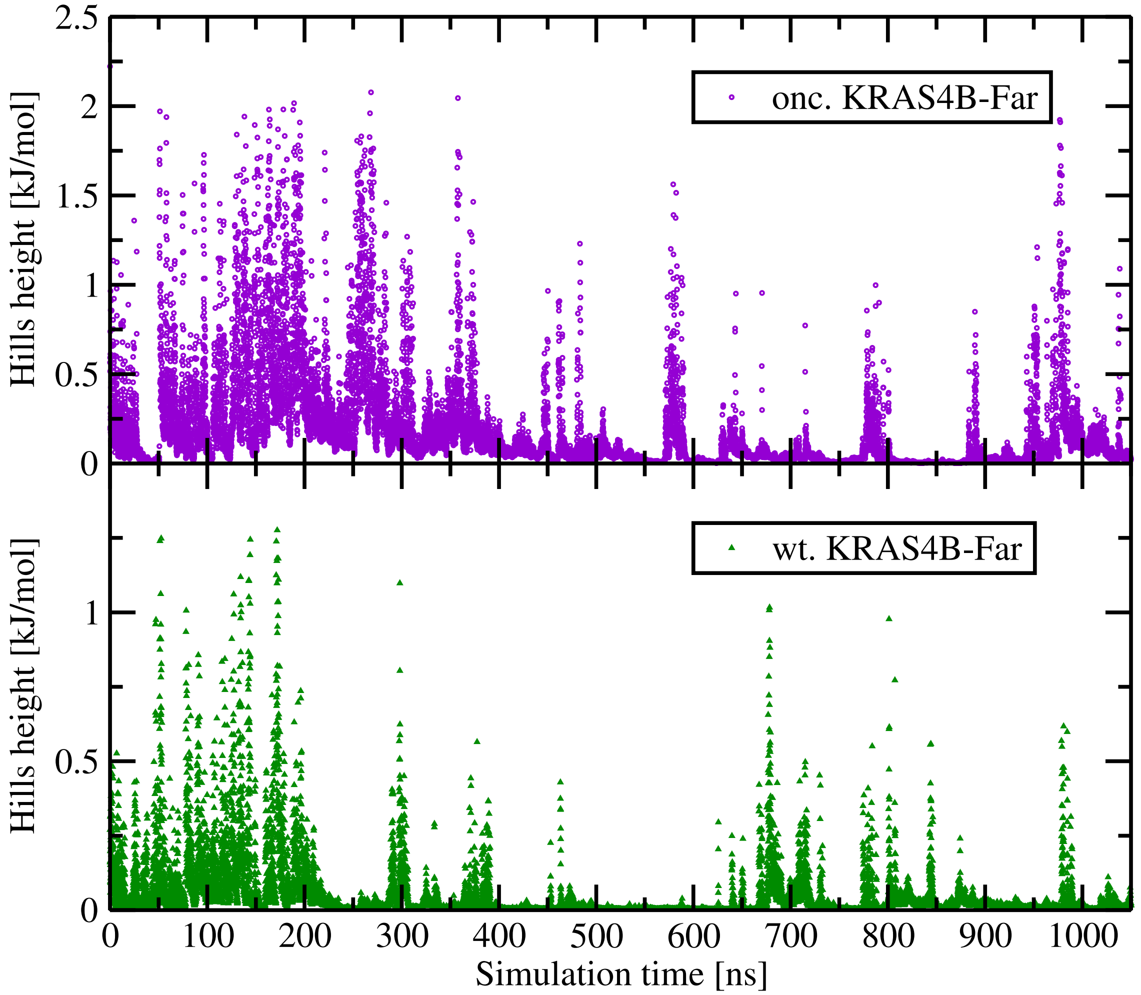


**Figure 10.** Well-tempered Metadynamics hills height as a function of time in different states of the farnesylated KRas-4B protein: oncogenic (top); wild-type (bottom).

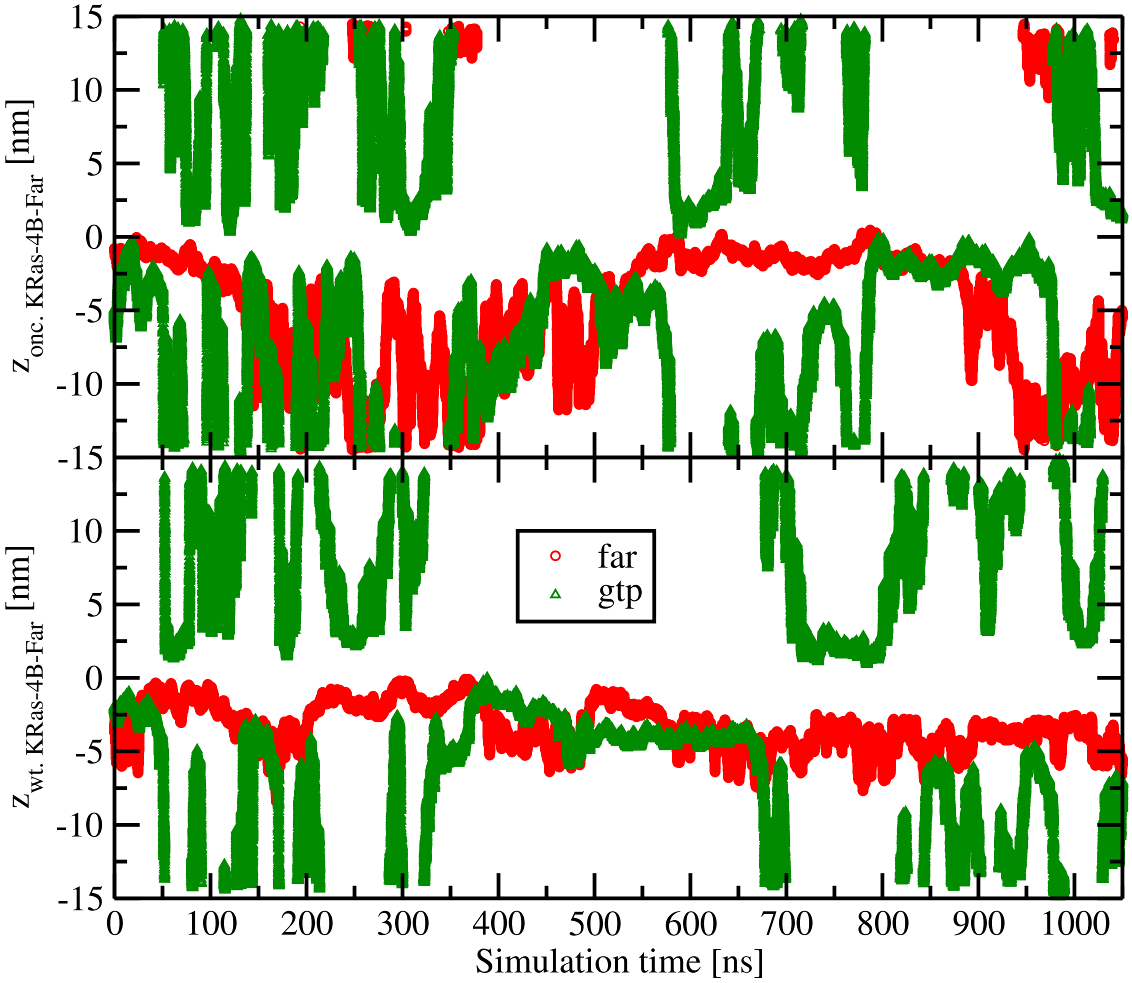


**Figure 11.** Numerical fluctuations of the collective variables as a function of time in different states of the farnesylated KRas-4B protein: oncogenic (top); wild-type (bottom).

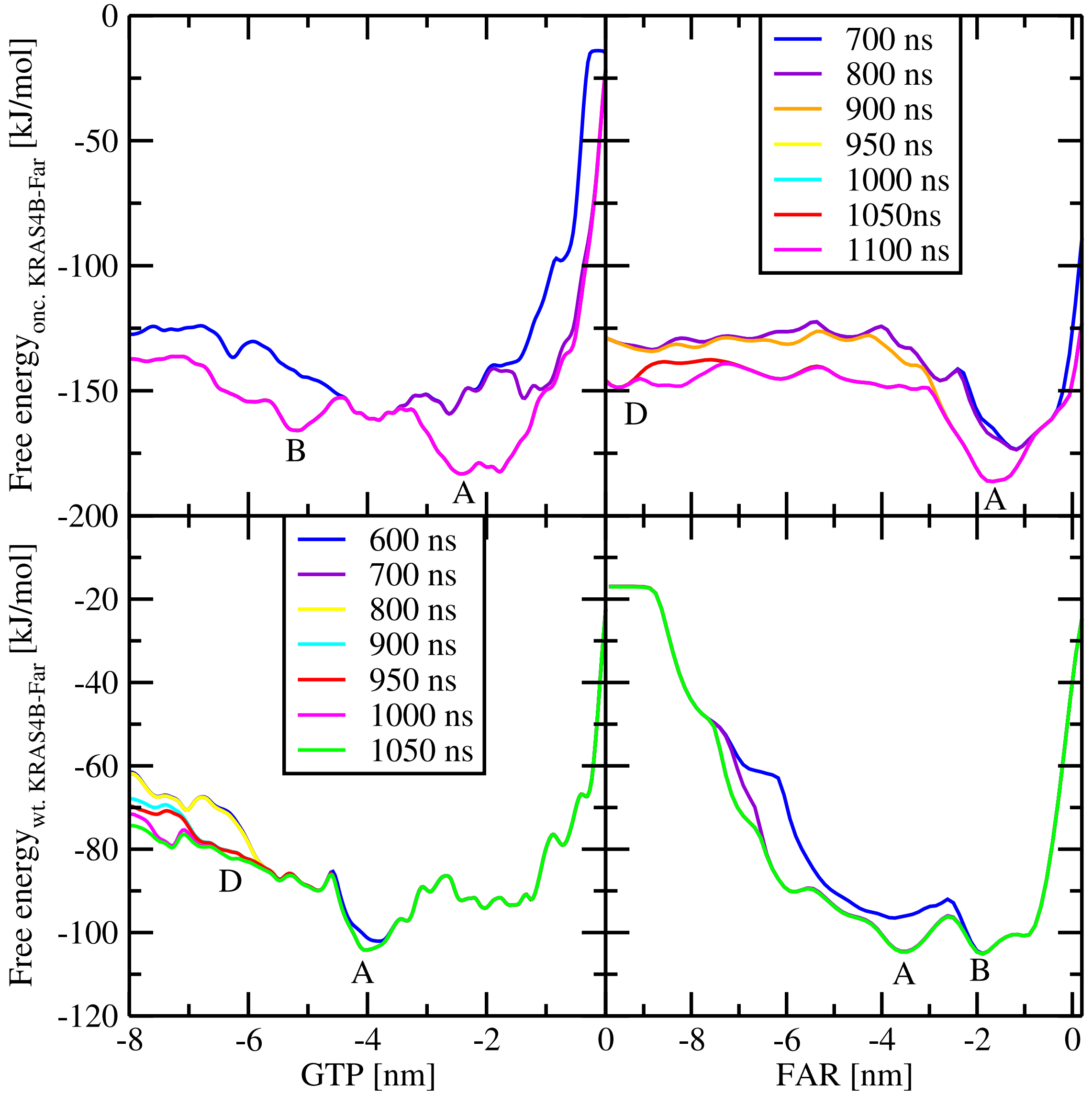


**Figure 12.** Time cumulative free energy profiles. Lower panels correspond to the wild-type system and upper panels to the oncogenic case. Labels 'A, B, D' correspond to relevant basins along the CV within the free energy profile with same labels as in Figs. 6 and 7 of the main text.

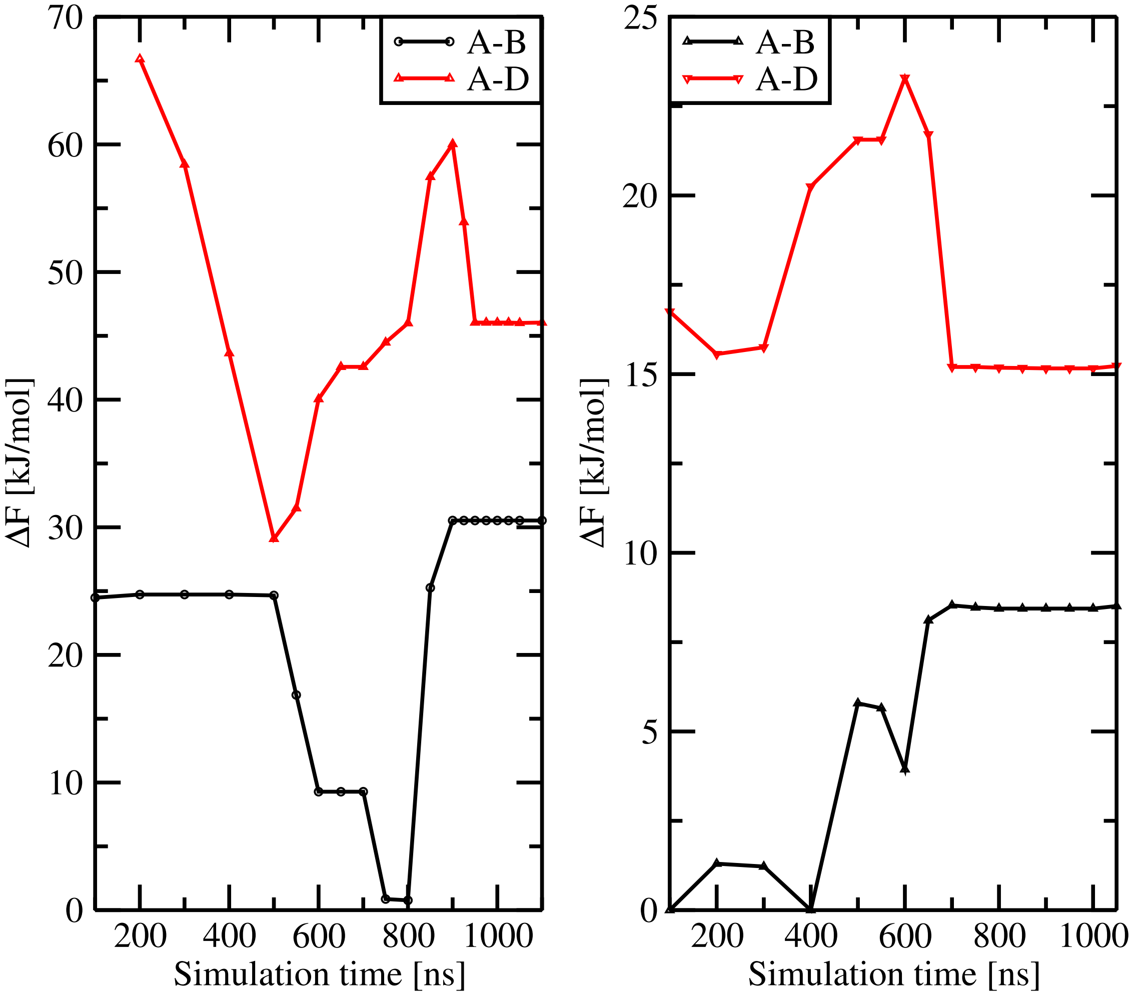


**Figure 13.** Convergence of well-tempered metadynamics based on the free energy barriers between two chosen basins. The selected basins have been shown in Fig. 12.
